# Aberrant splicing prediction during human organ development

**DOI:** 10.1101/2025.07.16.665183

**Authors:** Nils Wagner, Aleksandr M. Neverov, Alexandra C Martin-Geary, Shubhankar Londhe, Carina Schröder, Vicente A. Yépez, Nicola Whiffin, Julien Gagneur

**Affiliations:** School of Computation, Information and Technology, Technical University of Munich, Garching, Germany; Helmholtz Association - Munich School for Data Science (MUDS), Munich, Germany; Big Data Institute, University of Oxford, Oxford, UK; Centre for Human Genetics, University of Oxford, Oxford, UK; Broad Center for Mendelian Genomics, Program in Medical and Population Genetics, Broad Institute of MIT and Harvard, Cambridge, MA, USA; Institute of Human Genetics, School of Medicine and Health, Technical University of Munich, Munich, Germany; Computational Health Center, Helmholtz Munich, Neuherberg, Germany

## Abstract

Developmental disorders constitute a major class of genetic diseases, yet tools to identify splicing-disruptive variants during development are lacking. To address this need we extended the AbSplice framework to incorporate splicing dynamics across developmental stages. Moreover, we introduced several improvements including a continuous representation of splice site usage and integration of the rich set of predictions from the sequence-based model Pangolin. These advances improve the performance of the original model at predicting aberrant splicing events and help outperform purely sequence-based models including AlphaGenome, enabling predictions across embryonic, childhood, and adult tissues. Genome-wide scores for all single-nucleotide variants and a web interface to score indels are available to facilitate the exploration of the predictions. Our genome-wide predictions reveal a class of variants with splicing-disruptive effects confined to early development, particularly abundant in the brain (>18,000 variants). These variants are enriched in loss-of-function intolerant genes and neurodevelopmental disorder genes. Within the rare-disease cohort Solve-RD, high-impact brain-specific predictions are significantly enriched in individuals affected by neurodevelopmental disorders (NDD) in NDD-linked genes. Furthermore, the predicted developmental timing of the splicing disruption correlates with the clinical age of onset. Analysis of genomes of individuals with a suspected Mendelian disorder from Genomics England identified 26 unique variants in disease-linked genes, with stronger predicted effects during development than in adulthood, including a candidate new diagnosis in the gene *FGFR1*. Altogether, these results improve the accuracy of splice-disruptive variant prediction and provide tissue and developmental context to aid interpretation in rare disease diagnostics.

## Introduction

Aberrant splicing can disrupt gene function by altering RNA isoforms, leading to outcomes such as frameshifts, transcript degradation, or the loss of critical protein domains. If a variant strongly affects splicing isoform choice, the remaining abundance of functional RNA isoforms may be insufficient to maintain normal cellular function of the gene. Predicting such splicing aberrations is therefore critical for variant interpretation, particularly in the context of rare disease diagnostics and oncology^1–8^.

Aberrant splicing events are rare, large alterations of splice isoform usage and can be detected from RNA-seq data from human tissues^9–13^. Recently, we introduced AbSplice^14^, a model to predict the probability that a rare variant leads to aberrant splicing in a specific human tissue. AbSplice was trained on rare variants associated with aberrant splicing events for 49 adult tissues from the GTEx dataset^15^. AbSplice is intentionally lightweight: it does not retrain or replace sequence-based models, but contextualizes their predictions using empirical splice site usage. The model combines (i) tissue-agnostic sequence-based variant effect predictions from state-of-the art deep learning models with (ii) tissue-specific SpliceMaps derived from RNA-seq data quantifying splice site usage. Rather than modeling splicing de novo, the AbSplice framework can be understood as a contextual filter: it refines the context-agnostic sequence-based predictions by assessing whether a predicted variant effect is plausible and consequential in a specific context, based on empirical splice site usage. Notably, SpliceMaps can be generated from healthy and cancerous tissues^16^ and can be obtained from a handful of RNA-seq samples^14^, highlighting the framework’s adaptability. Predictions for new tissues or conditions do not require retraining the model but are obtained by supplying the corresponding SpliceMaps. Aberrant splicing predictions from AbSplice are valuable not only in isolation but also as input for downstream tasks such as aberrant gene expression prediction^17^ and rare variant association testing^18^.

The initial implementation of AbSplice used aberrant splicing events detected by FRASER^11^ as its ground truth for model training and evaluation. However, the more recently developed FRASER2^12^, based on a more robust intron-centric metric, offers a more accurate ground truth by reducing false-positive calls, enabling a more reliable assessment of model performance.

Advances in deep learning models for splicing prediction have further refined our ability to predict and interpret splicing alterations caused by variants (reviewed in ^19–21^). For instance, Pangolin^22^ shares the same context window and model architecture as SpliceAI^1^ yet demonstrated superior performance due to enhancements in both the quantity and quality of its training data. Specifically, Pangolin incorporates splicing quantifications derived from RNA-seq data across four tissues (heart, liver, brain, testis) and four mammalian species (human, rhesus macaque, mouse, rat). In contrast, SpliceAI was trained on binary labels of splice sites derived from the human standard genome annotation GENCODE^23^ as well as novel splice junctions commonly observed in the GTEx cohort. The initial implementation of AbSplice did not include Pangolin as the two tools were developed in parallel.

In parallel to this work, AlphaGenome^24^ was developed, which reports substantial improvements over other sequence-based models of splicing. Notably, when evaluating on tissue-specific aberrant splicing prediction, the authors adopted the AbSplice framework by combining AlphaGenome predictions together with tissue-specific SpliceMaps, effectively replacing the underlying sequence based scoring method in AbSplice. This shows that the AbSplice framework - the principle of integrating sequence-based predictions with empirical context from SpliceMaps - remains an effective strategy for aberrant splicing prediction and advocates for collecting a broader set of SpliceMaps.

During organ development, splicing events can undergo dynamic changes in inclusion levels, with brain and testis showing up to 20% of cassette exons being developmentally regulated^25^. Developmental alternative splicing events are highly conserved across evolution, underscoring their functional significance^25^. Variants impacting splicing at developmentally regulated sites may cause developmental disorders. However, their effects on splicing may be limited to specific stages, such as embryogenesis, and not manifest in adults (Fig. 1b). Since current splicing annotations, including the SpliceMaps^14^ used in the AbSplice model, are derived from adult RNA-seq data, the impact of variants on developmentally regulated splicing events is potentially underestimated. Thus, integrating developmental splicing dynamics into splicing prediction models should be considered to accurately assess variant effects across the full spectrum of an organism’s life stages.

**Fig. 1:**
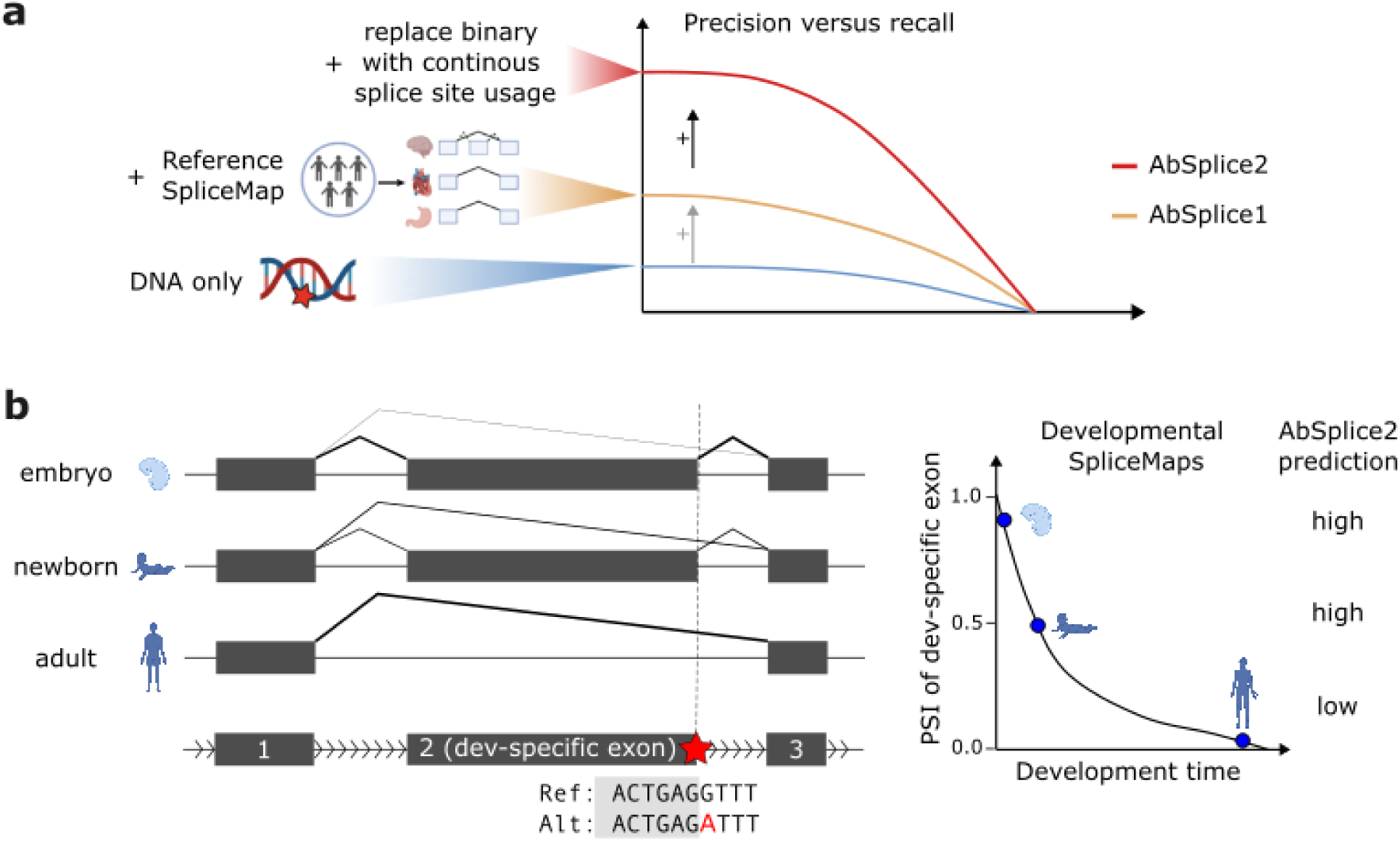
Improvements in the AbSplice framework and extension to variant effect prediction around developmentally regulated splice sites. **a)** We evaluated the performance of splicing prediction tools on GTEx splicing outliers^11^. AbSplice1 already improved sequence-only predictors by leveraging tissue-specific SpliceMaps. Replacing binary estimates of splice site usage with a continuous metric led to a further two-fold precision increase. **b)** Variants (red star) in the vicinity of developmentally regulated alternative splice sites exhibit changing effects over the course of organ development. These dynamic changes are captured by developmental SpliceMaps and reflected in AbSplice2 predictions.

Here, we present an updated version of AbSplice (referred to as AbSplice2) that addresses these issues. We evaluate models on the improved ground truth from FRASER2 and integrate state-of-the-art splicing predictions from Pangolin (Fig. 1a). Furthermore, we extend our analyses to include developmental splicing dynamics (Fig. 1b). Together, this led to improved detection of aberrant splicing causing variants while providing developmental and tissue-specific context.

## Material and methods

### Datasets

#### GTEx

We downloaded the RNA-seq read alignment files (BAM files) and the variant calling files (VCF files) from WGS from GTEx v8p (hg38) from the database of Genotypes and Phenotypes (dbGaP) (study accession: phs000424.v8.p2). We used data from 946 individuals with paired WGS and RNA-seq measurements (n = 16,213) in at least one tissue.

#### BrainGVEx

We downloaded the RNA-seq read alignment files (BAM files) and the variant calling files (VCF files) from WGS of the BrainGVEx dataset from the PsychENCODE portal (SynID: syn4590909^26^). We used a total of 230 samples isolated from the Brain Frontal Cortex of individuals with bipolar disorder or schizophrenia as well as from neurotypical controls.

### Developmental dataset

Annotations for devAS events were taken from Supplemental files of ^25^. The RNA-seq data for all developmental stages and tissues was downloaded from^27^.

### Data preprocessing

Most data preprocessing was done analogously to our previous publication^14^. We describe here only the differences, namely replacing our ground truth of aberrantly spliced events called by FRASER with events called by FRASER2.

#### Splicing outlier detection

FRASER2 (v1.99.4) was run in the different datasets via the Detection of RNA Outliers Pipeline^28^ (DROP, v.1.4.0) with the default parameters and cut-offs. The main difference between FRASER2 and FRASER is the use of a new intron-specific metric (called intron Jaccard index) replacing the previous splice-site specific metrics. Significant junctions were those with an absolute splice deviation (delta Jaccard) > 0.1 and a FDR < 0.1. We additionally applied the rare variant filter as described in^14^, filtering for splicing outliers containing a rare variant in the vicinity of ±250 bp of every splice site based on RNA-seq from the sample. Importantly, this filter was applied to all splice sites identified by FRASER2, which includes both annotated splice sites as well as cryptic splice sites.

### Aberrant splicing prediction benchmark

The benchmark was done analogously to our previous study^14^, with the task being to predict whether a protein coding gene with one or more rare variants within the gene body is aberrantly spliced in a given tissue of an individual. The performance of models was evaluated using precision-recall curves and the average precision score.

### Tissue-specific SpliceMap

SpliceMaps were created analogously to our previous study^14^. In short, SpliceMaps list active introns together with aggregate metrics quantifying splicing usage. These aggregate metrics are calculated separately for donor and acceptor sites. In the context of AbSplice the relevant metrics are the median number of split reads (across samples used to create the SpliceMap) sharing the same splice site as well as the reference isoform proportion of the intron, which is used to translate the predicted tissue-agnostic variant effect of MMSplice^2^ to a given tissue using the splicing scaling law^14,29^.

#### Developmental SpliceMaps

We created developmental SpliceMaps by using RNA-seq data from^27^ in 7 human tissues (brain, cerebellum, heart, liver, testis, ovary, kidney) and for 15 different developmental timepoints (from 4 weeks post conception to adult stages). Due to small cohort sizes, each SpliceMap for a given developmental stage was created from samples corresponding to the same tissue using a sliding window of size 1 for the developmental stage.

### Comparison of SpliceMaps with SpliceVault

To assess the coverage of splice sites detected in the GTEx and developmental SpliceMaps with other population-scale splicing catalogs, we compared splice site positions across four sources: GTEx SpliceMaps (49 tissues^14^), developmental SpliceMaps (7 tissues across 15 developmental timepoints, this study), the SpliceVault 300K database^30^, and the GENCODE v38 annotation. All analyses were restricted to protein-coding genes.

All splice sites from SpliceMaps were included, including rare sites detectable only in a small subset of RNA-seq samples of a given tissue. To ensure a direct comparison with SpliceVault’s inclusive methodology, we additionally generated an “unfiltered” version of the GTEx SpliceMaps, omitting the standard reproducibility filter – which normally requires an intron to be supported by at least one split-read in more than 5% of samples within a tissue. This allows for a direct comparison with the SpliceVault database, which catalogs all detectable stochastic splicing events regardless of population frequency. For developmental SpliceMaps, where only 1 to 5 samples were available per tissue-time point pair, we defined “replicated” as being present in at least 2 samples instead of the 5% reproducibility filter. SpliceMap coordinates follow the BED-file convention (0-based start, 1-based end); a positional correction of +1 was applied to start coordinates to match the 1-based coordinate system used by SpliceVault. Genomic coordinates in developmental SpliceMaps were lifted from hg19 to hg38 using pyliftover^31^.

For SpliceVault, we utilized the 300K-RNA repository, which collates unannotated splicing events across 335,663 samples. We further partitioned these data into SRA (n=316,449) and GTEx (n=19,214) sources.

### Aberrant splicing prediction models

Some aberrant splicing prediction models (MMSplice, AbSplice1) were used analogously to our previous study^14^. We highlight here only the changes.

#### Pangolin

Pangolin is a deep learning model that predicts the effect of a variant on the splice site usage. It computes a gain and a loss score for every position within a user-defined window around the variant that represents the increase and decrease in the usage of a potential splice site at the respective positions. Pangolin outputs the maximum gain and the maximum loss scores within the window together with the corresponding positions. It also provides an option to mask scores when a genome annotation is provided to the model, which sets those scores to zero if Pangolin predicts activation for annotated splice sites and deactivation for unannotated splice sites. We ran Pangolin with the default settings, i.e. a window of length 50 nt and masking using the Gencode annotation (release 38). We computed genome-wide predictions for all possible SNVs in protein coding genes which are made publicly available (https://doi.org/10.5281/zenodo.15649338).

We intersected the genomic coordinates of the predicted affected splice site position with splice sites annotated in the tissue-specific SpliceMaps. The splice site usage of the affected splice site along with the variant effect score were provided as input features to the AbSplice2 model.

#### AbSplice2

Training of the AbSplice2 model was analogous to our previous study^14^ changing the input features of the model and using FRASER2 outlier labels as a ground truth. While the recommended delta Jaccard cutoff in FRASER2 is 0.1, the reported AbSplice2 model was trained using a more stringent cutoff of 0.3 due to a discrepancy in the processing pipeline that was identified during the revision process of this manuscript. We confirmed that retraining AbSplice2 with the optimal 0.1 cutoff had only a marginal impact on overall performance (Average Precision = 0.27 vs 0.28) and yielded highly correlated prediction probabilities (Pearson correlation = 0.96). As the choice of training threshold has a negligible impact on the learned parameters, the originally deployed model was retained. The features of the final AbSplice2 model were the prediction score from MMSplice + SpliceMap, MMSplice + SpliceMap + ψ_ref, the predicted loss and gain scores from Pangolin and a continuous feature from SpliceMap quantifying splice site usage in the target tissue (the median number of split reads per sample sharing the same acceptor/donor site, depending on which splice junction was used to score the variant in MMSplice + SpliceMap) as well as the splice site usage of the predicted affected site by Pangolin.

The reported performance on the aberrant splicing prediction benchmark (GTEx) is based on fivefold-stratified cross-validation, grouped by individuals to prevent information leakage. The final model which was used for genome-wide predictions and validations in independent datasets was trained on the full GTEx dataset.

We computed genome-wide predictions for all possible SNVs in protein coding genes which are made publicly available (https://doi.org/10.5281/zenodo.15644548).

#### Feature contributions to model performance

To quantify the marginal contribution of each component improvement to AbSplice2 performance, we constructed pairwise matched comparisons across four feature dimensions – base model choice (SpliceAI vs. Pangolin), SpliceMap integration (no SpliceMap vs. SpliceMap using binary splice site expression threshold), signed features (unsigned maximum score vs. directional signed scores), and continuous splice site usage (binary threshold vs. continuous median split-read count) – such that each pair of models differed in exactly one dimension. For each matched pair and each of the 49 GTEx tissues, we computed the marginal gain in auPRC as the difference between the two models. The average marginal gain attributable to a given feature dimension was computed as the mean across all matched pairs sharing that dimension. Median marginal gains and interquartile ranges (25th–75th percentile) were summarized across tissues.

### Comparisons with AlphaGenome, Nucleotide Transformer v3 and SpliceTransformer

Due to computational constraints, we could not predict variant effects for the full GTEx aberrant splicing benchmark dataset (8.1 million rare variants in total). We constructed a model-informed benchmark subset consisting of: (i) all variants linked to either FRASER1 or FRASER2 splicing outliers (n = 14,647); (ii) non-outlier variants ranked above conservative model thresholds by at least one of the available splicing models (AbSplice1-DNA ≥ 0.01, AbSplice2-DNA ≥ 0.01, SpliceAI Delta Score ≥ 0.1, |Pangolin gain score| ≥ 0.1, |Pangolin loss score| ≥ 0.1, or |ΔPSI of MMsplice with SpliceMap| ≥ 0.1; n = 113,086); and (iii) non-outlier variants below all of these thresholds (n = 60,000; randomly sampled). This subset consisted of 187,733 unique variants. Because all outlier variants are included, performance on this subset is conservative relative to the full benchmark: additional variants from the full benchmark can only contribute false positives, not additional true positives. All performance evaluations of AlphaGenome, Nucleotide Transformer v3, and SpliceTransformer are based on this subset.

#### AlphaGenome

Variants were scored using the AlphaGenome API (version 0.5.1) with a 1 Mb sequence context window. Using the recommended variant scorers, we obtained (i) tissue-agnostic splice site scores (SPLICE_SITES), (ii) tissue-specific splice site usage scores (SPLICE_SITE_USAGE), and (iii) tissue-specific splice junction scores (SPLICE_JUNCTIONS). Scores were aggregated per variant–gene–tissue by taking the maximum absolute score across all predicted positions within the gene. We computed a composite splicing score following the AlphaGenome paper: composite = SPLICE_SITES + SPLICE_SITE_USAGE + (SPLICE_JUNCTIONS / 5).

#### AbSplice models using AlphaGenome

AbSplice models using AlphaGenome scores as base model inputs (referred to as AbSplice-AG models) were trained and evaluated on the benchmark subset using stratified five-fold cross-validation on individuals. For each of the four AlphaGenome score types (sites, junctions, usage and the composite score), we trained models using either the tissue-specific score or a tissue-agnostic proxy (maximum absolute value across GTEx tissues per variant–gene pair) as the base model feature, combined with either a binary splice site expression indicator (as in AbSplice1) or the continuous median number of split reads (as in AbSplice2) as the SpliceMap feature.

The best AbSplice-AG model combined five AlphaGenome score types – splice site, splice junction, splice site usage, RNA activity and RNA log-fold change – with the continuous median split-read coverage as the SpliceMap feature. Performance was compared to an AbSplice2 model that was retrained on the same subset. To assess the contribution of directional features to AbSplice2’s performance, we additionally trained a model without directional features, replacing the directional Pangolin gain and loss scores with the absolute maximum score over those, and the signed MMSplice delta logit PSI and delta PSI with their absolute values.

#### Nucleotide Transformer v3

Nucleotide Transformer v3 (NTv3) is a genomic foundation model pre-trained on ∼9 trillion base pairs spanning more than 128,000 species and post-trained on ∼16,000 functional genomic tracks^32^. Unlike AlphaGenome, NTv3 does not provide a recommended scoring scheme for splicing variant effect prediction. We therefore implemented our own approach based on the output tracks of the post-trained model. Variants were scored using the post-trained NTv3 100M model (NTv3_100M_post) with a 32 kb sequence window centered on the variant position (hg38 reference). Reference and alternate sequences were scored independently, and variant effects were computed as *alt - ref* differences in the predicted output tracks. We derived two scores. A tissue-agnostic splice-site score was computed as the maximum of splice donor and splice acceptor gain (and loss) scores within a ±50 bp window around the variant, where gain (loss) is the maximum positive (negative) change in the BED element class probabilities for splice donor or acceptor. A tissue-specific score was derived: at the position of maximum splice donor or acceptor gain or loss, we computed the delta of the tissue-averaged RNA-seq bigwig signal. The tissue-specific score is the maximum absolute bigwig delta across the four splice-site event positions (donor gain, donor loss, acceptor gain, acceptor loss). Tissue-specific scores were available for 34 of the 49 GTEx tissues used in the benchmark. GTEx tissues that did not have a NTv3 prediction available were dropped in the performance analysis.

#### SpliceTransformer

Variant effect scores for SpliceTransformer were obtained by following the instructions in the publicly available SpliceTransformer GitHub repository^33^. SpliceTransformer provides tissue-specific predictions for 15 meta-tissues (e.g. *Brain*). In the benchmark we used the same score for each GTEx tissue corresponding to the meta-tissue (e.g. SpliceTransformer’s *Brain* prediction used for all 13 of GTEx *Brain* tissues: e.g. *Brain Frontal Cortex, Brain Hypothalamus*), resulting in 33 tissue matches. GTEx tissues that did not have a match were dropped in the performance analysis of SpliceTransformer.

### Developmental variants genome-wide

We computed all possible SNVs within a 100 bp window from all splice sites related to devAS events annotated in the Supplemental files from Mazin et al.^25^. We used SpliceMaps from all 15 developmental stages and all 7 tissues to compute AbSplice2 scores for each developmental stage.

We defined a variant to have a *developmental* effect in a tissue, if it had an AbSplice score above the high cutoff (0.2) at any timepoint during embryogenesis up to toddler age. A variant was defined to have a *purely developmental* effect if it had additionally a low score (below 0.05) in the corresponding adult GTEx tissue. Similarly, we defined a variant to have an *adult* effect in a tissue if it had an AbSplice score above the high cutoff (0.2) in GTEx. A variant was defined to have a *purely adult* effect, if it had additionally a low score (below 0.05) at all timepoints during embryogenesis up to toddler age in the corresponding tissue. For tissues with several subtissues in GTEx (e.g. brain), the maximum over all subtissues was taken.

### Analysis of variants in Solve-RD

We computed variant effect scores for all rare variants in 24,560 individuals within the Solve-RD cohort, consisting of 15,028 affected and 9,532 unaffected individuals. We defined a rare variant to have a minor allele frequency in gnomAD exomes and genomes below 0.1%. We grouped affected individuals into early (up to the age of 5 years, N=2,378) and late age of onset (after 40 years of age, N=502). Among the affected individuals, 4,780 were diagnosed with a neurological or neurodevelopmental disorder, of which 1,033 belonged to the early age of onset group and 365 belonged to the late age of onset group.

The disease gene list for neurological and neurodevelopmental disorders (n=1,616) was downloaded from GeneTrek^34^, using the column ‘SysNDD definitive genes (updated 2023-11-09)’.

Enrichments of high impact variant effect predictions were computed using the odds ratio, measuring the odds of observing a high-impact variant in a test group compared to a reference group. For the enrichment analysis in Fig. S6b, the test group consisted of variants in all affected individuals, and the reference group consisted of variants in all unaffected individuals. For the enrichment analysis in Fig. 2e, the test group consisted of variants in individuals affected with neurological or neurodevelopmental disorders and within NDD-related genes. For the age-of-onset analysis (Fig. 4d), we restricted the calculation to affected individuals whose disease onset was categorized as either ’early’ (embryogenesis up to age 5 years) or ‘adult’ (after 40 years of age). The odds ratio for *purely developmental*, *developmental*, *adult* and *purely adult* variants was computed based on these two extreme groups.

**Fig. 2:**
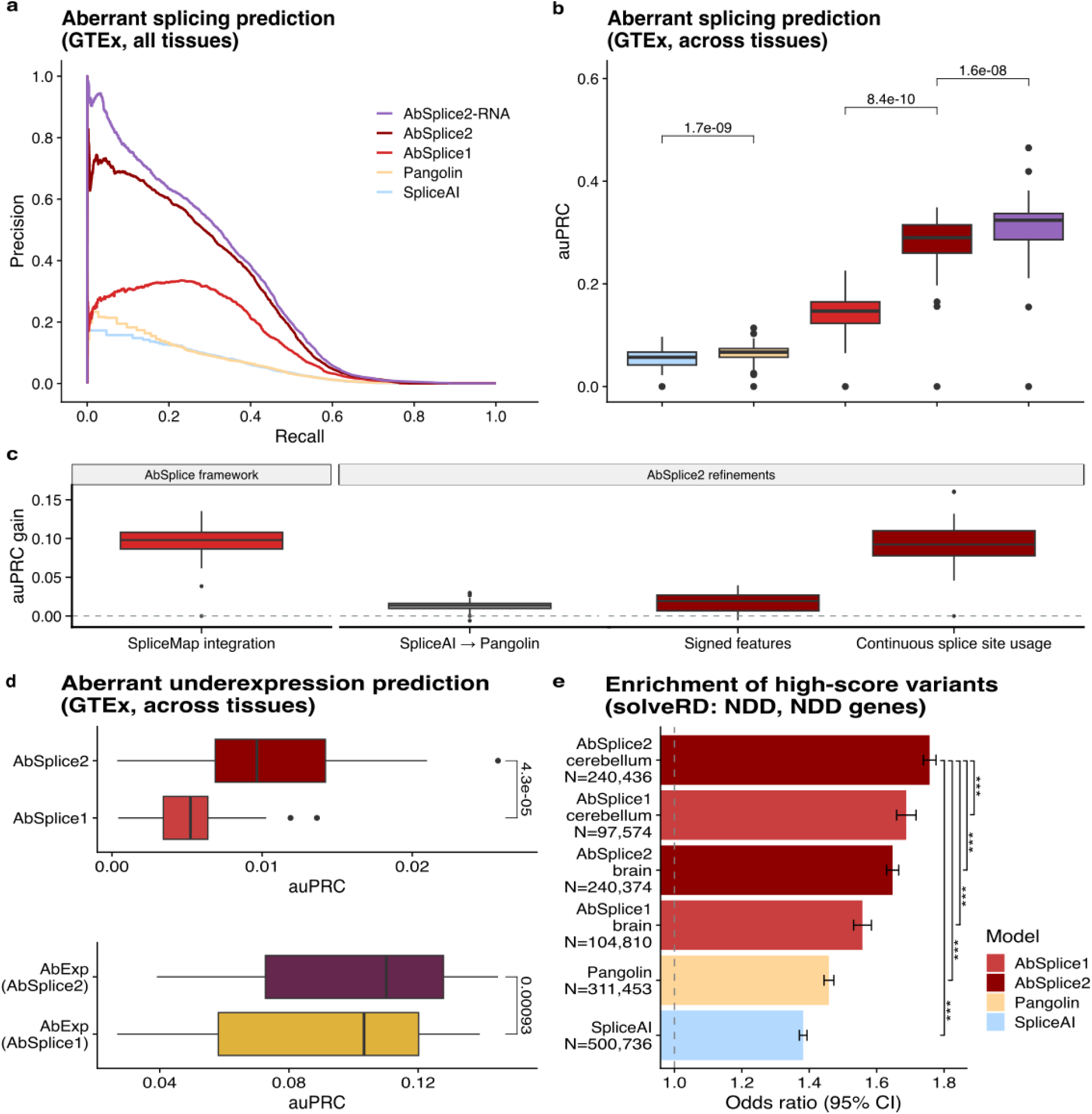
Improvements for prediction of aberrant splicing and underexpression. **a)** Precision–recall curve comparing the overall prediction performance on all GTEx tissues of the models SpliceAI, Pangolin, AbSplice1, and AbSplice2. All models are evaluated on FRASER2 outlier calls. The color legend in (**a**) is shared across (**a**-**b**). **b)** Distribution of the area under the precision–recall curve (auPRC) of the models in (**a**) across tissues (n = 49). Center line, median; box limits, first and third quartiles; whiskers span all data within 1.5 interquartile ranges of the lower and upper quartiles. P values were computed using the paired one-sided Wilcoxon test. **c)** Marginal contribution of AbSplice components to prediction performance. Boxes show the distribution of average marginal auPRC gains across 49 GTEx tissues attributable to each AbSplice design component, organized into two groups: AbSplice framework (SpliceMap integration: the core AbSplice approach that converts raw sequence predictions of a tissue-agnostic base model into tissue-aware aberrant splicing scores by integrating tissue-specific splicing data) and AbSplice2 refinements. AbSplice2 refinements: base model (replacing SpliceAI with Pangolin as the underlying sequence-based variant effect predictor), signed features (using directional gain/loss scores instead of the unsigned maximum) and continuous splice site usage (replacing a binary ≥10 split-read expression threshold with the continuous median split-read count per splice site). For each component, the marginal gain per tissue was computed as the average over all matched model pairs that differ only in that dimension, holding all other features constant. Center line: median; box limits: first and third quartiles; whiskers extend to 1.5× the interquartile range. The dashed line indicates zero gain. **d)** Distribution of the auPRC for AbSplice1 and AbSplice2 as well as the aberrant gene underexpression prediction model AbExp (using AbSplice1 or AbSplice2 as input features) on the task of predicting aberrant gene underexpression in the GTEx dataset. **e)** Enrichment of high impact variants (SpliceAI: 0.8, Pangolin: 0.8, AbSplice1: 0.2, AbSplice2: 0.2) in affected individuals diagnosed with neurological and neurodevelopmental disorders within the Solve-RD cohort. The odds ratios are computed across neurodevelopmental disorder disease genes. Note that AbSplice predictions for brain and cerebellum show the largest enrichments. Error bars indicate Wald 95% confidence intervals. Asterisks mark significance levels of Wald z-tests on log odds ratios, comparing each model to the AbSplice2 cerebellum prediction (*<0.05, **<0.01, ***<0.001).

### Analysis of variants in the NGRL

We identified 39,915 participants from the Genomics England 100,000 Genomes Project (100k) and 24,821 participants from the Genomic Medicine Service (GMS) datasets within the UK NGRL. Consenting probands were selected where they had a genome sequence obtained from blood samples, aligned to GRCh38, and where variants were called using the ‘V4’ pipeline.

Where multiple alignment files existed, the most recent was selected, and for any remaining duplicates, the specific variant file used in the GEL interpretation pipeline was chosen. We selected all variants within 100bp of devAS splice sites. In total this resulted in n=1,820 unique variants in 100k and n=1,135 unique variants in GMS, including both common and rare variants.

Candidate variants were filtered to those that were fully genotyped and passed GEL’s ‘Stringent’^35^ filters (n=552 unique variants in 100k, 272 unique variants in GMS). Variants were further filtered to those that are in ‘Green’ PanelApp genes for each participant’s phenotype (n=22 unique variants in 100k, and 17 unique variants in GMS).

We further subset variants to those where the AbSplice2 score (in any of the seven tissues with developmental data) was greater at early developmental stages compared to the adult prediction (n= 15 unique variants in 100k, 13 unique variants in GMS). 26 unique variants in 47 individuals were represented in total.

One variant (chr8:67203252 C/T, NM_006421.5:c.4960-1G>A) was identified in a ‘solved’ case in the NGRL Exit Questionnaire table, with this variant provided as the diagnostic variant (Table 1) (positive controls). Eight additional participants were flagged as ‘solved’, with an alternative variant recorded. We compared these participants’ Human Phenotype Ontology (HPO) terms with the HPO terms associated with both the gene in the exit questionnaire, and the gene identified using AbSplice2. None were supportive of an expansion to the existing diagnosis.

**Table 1:**
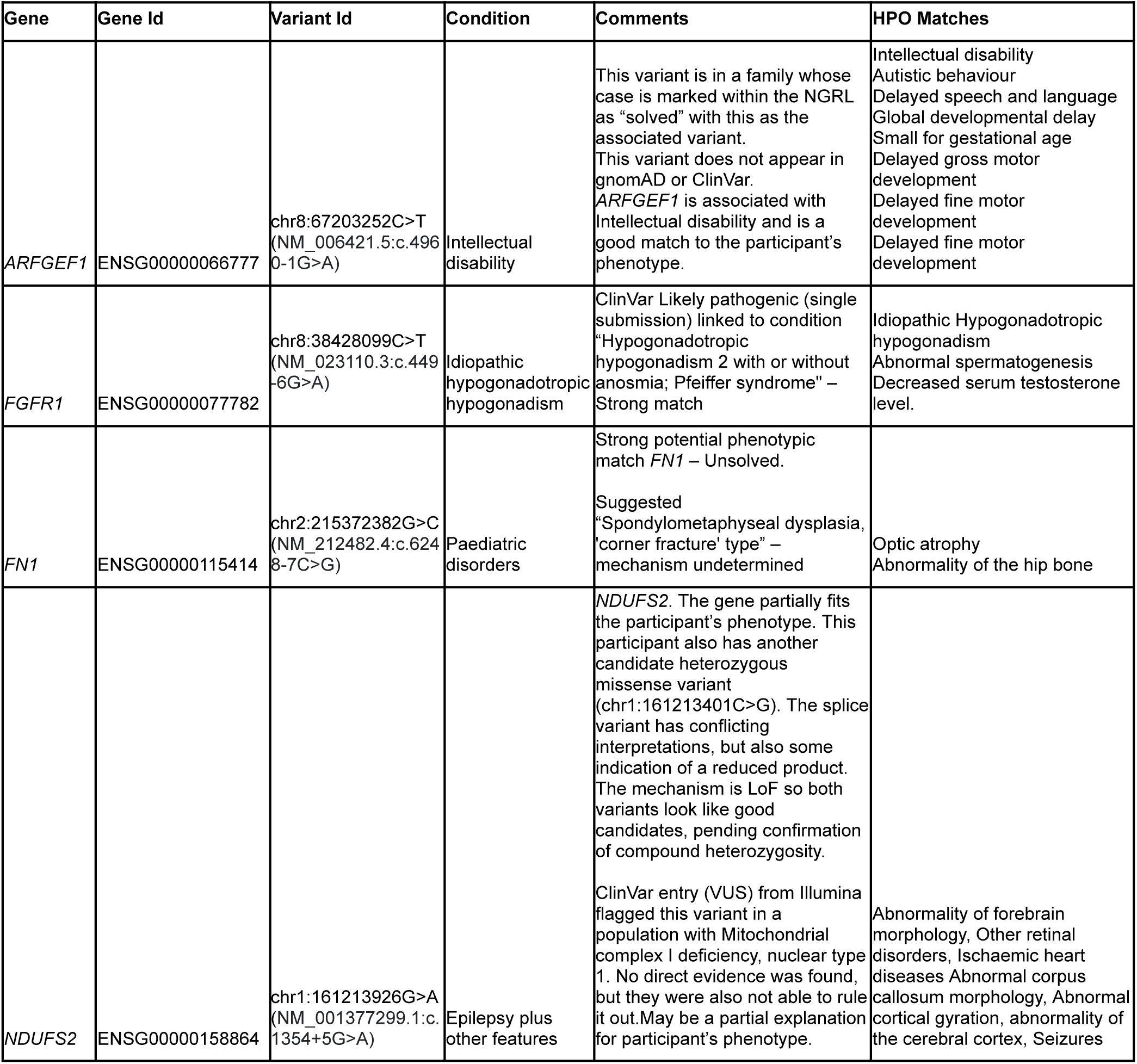
Prioritized variants by applying AbSplice2 on the NGRL rare disorder cohorts. HPO matches: overlapping Human Phenotype Ontology terms between the participant and the gene associated with the AbSplice2 candidate variant. Variant coordinates are in hg38.

| Gene | Gene Id | Variant Id | Condition | Comments | HPO Matches |
|---|---|---|---|---|---|
| *ARFGEF1* | ENSG00000066777 | chr8:67203252C>T<br>(NM_006421.5:c.4960-1G>A) | Intellectual disability | This variant is in a family whose case is marked within the NGRL as “solved” with this as the associated variant.<br>This variant does not appear in gnomAD or ClinVar.<br>*ARFGEF1* is associated with Intellectual disability and is a good match to the participant’s phenotype. | Intellectual disability<br>Autistic behaviour<br>Delayed speech and language<br>Global developmental delay<br>Small for gestational age<br>Delayed gross motor development<br>Delayed fine motor development<br>Delayed fine motor development |
| *FGFR1* | ENSG00000077782 | chr8:38428099C>T<br>(NM_023110.3:c.449-6G>A) | Idiopathic hypogonadotropic hypogonadism | ClinVar Likely pathogenic (single submission) linked to condition “Hypogonadotropic hypogonadism 2 with or without anosmia; Pfeiffer syndrome" – Strong match | Idiopathic Hypogonadotropic hypogonadism<br>Abnormal spermatogenesis<br>Decreased serum testosterone level. |
| *FN1* | ENSG00000115414 | chr2:215372382G>C<br>(NM_212482.4:c.6248-7C>G) | Paediatric disorders | Strong potential phenotypic match *FN1* – Unsolved.<br>Suggested “Spondylometaphyseal dysplasia, 'corner fracture' type” – mechanism undetermined | Optic atrophy<br>Abnormality of the hip bone |
| *NDUFS2* | ENSG00000158864 | chr1:161213926G>A<br>(NM_001377299.1:c.1354+5G>A) | Epilepsy plus other features | *NDUFS2*. The gene partially fits the participant’s phenotype. This participant also has another candidate heterozygous missense variant (chr1:161213401C>G). The splice variant has conflicting interpretations, but also some indication of a reduced product. The mechanism is LoF so both variants look like good candidates, pending confirmation of compound heterozygosity.<br>ClinVar entry (VUS) from Illumina flagged this variant in a population with Mitochondrial complex I deficiency, nuclear type 1. No direct evidence was found, but they were also not able to rule it out.May be a partial explanation for participant’s phenotype. | Abnormality of forebrain morphology, Other retinal disorders, Ischaemic heart diseases Abnormal corpus callosum morphology, Abnormal cortical gyration, abnormality of the cerebral cortex, Seizures |

Each potential novel diagnostic variant was then assessed based on:

- Allele frequency from gnomAD

- Presence in ClinVar - removed if benign, no conflicts

- The genotype of the variant matched the reported gene-disease mechanism in DECIPHER

- Phenotypic overlap was assessed by comparing HPO terms provided for each participant, with HPO terms associated with the gene in the Human Phenotype Ontology^36^, accounting for frequency of association where available.

For plausible variants in genes with likely biallelic inheritance (n=23), we searched for a clear second pathogenic variant within the MANE transcript coordinates. Any additional variants were passed to Ensembl’s VEP and assessed based on gnomAD (AF < 0.05), predicted consequence (missense, splice region, frameshift, stop-gained, start-lost) and LOFTEE high confidence LoF annotation (in line with known mechanisms for that gene as defined in DECIPHER), SpliceAI score > 0.2 CADD (> 25.3) and PhyloP (> 7.367) scores, and if present in ClinVar that they had not been classified as benign).

After this assessment, we prioritised three candidate variants (Table 1). For the purpose of anonymisation any combination of HPO terms for a given participant that were shared by fewer than 5 participants in the 100,000 genomes project were adjusted to parent/grandparent HPO terms where necessary, to ensure participants were not identifiable.

### AbSplice web interface

A web interface (https://absplice.cmm.cit.tum.de) allows users to run AbSplice on input SNVs and indels. As input, it requires a list of up to 10 variants and the genome build (hg19 or hg38). In addition, the user can select the tissue(s) of interest out of 49 GTEx tissues (the default is all) as well as tissues corresponding to 15 different developmental timepoints (spanning embryogenesis to adult stages) in brain, cerebellum, heart, kidney, liver, ovary and testis. The output displays the variant, Ensembl gene ID, HGNC Symbol, tissue, and AbSplice2 score, as well as Pangolin and MMSplice scores. The output can be downloaded as a CSV file.

## Results

### Contribution of AbSplice2 design choices to model performance

AbSplice was originally trained and evaluated using the aberrant splicing caller FRASER applied to the Genotype-Tissue Expression (GTEx) dataset^11,14^. FRASER2, a refined version of FRASER, was shown to reduce false positives in outlier detection by about one order of magnitude at minimal loss of sensitivity^12^. We detected 1,462,199 outlier introns with FRASER and 254,210 with FRASER2 across GTEx tissues (5.8 times fewer). The AbSplice pipeline^14^ had already incorporated a rare-variant-based post-filter to suppress likely false positive outliers from the original FRASER, explaining why the remaining difference between the two ground truths after filtering was more modest with 23,390 outlier events for FRASER and 21,482 for FRASER2.

Since no RNA-seq-based outlier caller provides a perfect ground truth, we assessed how model performance depends on the choice of evaluation framework. Evaluating all models separately within each ground truth (Fig. S1a), we found that the relative ranking of models was preserved: AbSplice consistently outperformed SpliceAI and Pangolin regardless of which ground truth was used for evaluation. To assess how the ground truth affects model training, we retrained AbSplice on FRASER2 and cross-evaluated the models. We found that while training and evaluating on the same ground truth source performed marginally, yet significantly, better than using a different ground truth source for training, the overall impact on performance was small (Fig. S1a). Furthermore, retraining had only a minor effect on the learned feature contributions (Fig. S1b), demonstrating that the framework is robust to imperfect ground truth data and primarily captures the underlying biological signal rather than caller-specific biases. We therefore adopted FRASER2 as the ground truth throughout this work.

AbSplice1 was based on variant effect predictions of the tissue-agnostic deep learning models MMSplice^2^ and SpliceAI^1^ which were combined with tissue-specific splicing annotations (SpliceMaps)^14^. In developing AbSplice2, we introduced three design changes and systematically quantified their individual contributions to prediction performance through pairwise matched comparisons across 49 GTEx tissues (Fig. 2c, Methods).

Pangolin is a deep learning model that uses a similar model architecture as SpliceAI. While SpliceAI was purely trained on genomic annotations, Pangolin additionally used quantitative usage levels of splicing in 4 different tissues and was trained across 4 mammalian species. Despite this expanded training, replacing SpliceAI with Pangolin contributed a modest median gain of +0.015 auPRC per tissue (IQR: +0.010–0.019; P = 1.3×10⁻⁸), in line with published benchmarks showing consistent, but small differences between these two sequence models^22,37,38^.

Both SpliceAI and Pangolin provide separate scores for splice site usage increase and decrease. In AbSplice1, we used only the absolute maximum variant effect score (unsigned) of SpliceAI. However, the direction of change is also informative: using directional (signed) scores enables the model to learn interactions with other signed features such as MMSplice predictions, providing a more reliable basis for predictions when models agree on the direction of change. Replacing the unsigned maximum with the predicted splice site gain and loss scores improved performance by a median of +0.027 auPRC per tissue (IQR: +0.011–0.039; P = 3.3×10⁻⁶).

To discriminate between genuine splicing events and splicing noise, AbSplice1 employed a binary threshold on the median number of split reads supporting a splice site across samples capturing splice site usage. However, thresholding can be highly sensitive to small variations in the data near the threshold and lead to drastically distinct predictions and misleading interpretations. By learning a continuous sigmoid-shaped effect of splice site usage instead of a binary threshold, we achieved the largest marginal improvement among the three components: a median gain of +0.087 auPRC per tissue (IQR: +0.069–0.108; P = 1.8×10⁻¹⁵).. This adjustment reduced abrupt prediction shifts due to minor fluctuations in splice site usage, enhancing model reliability. The contribution scores on the plateaus were similar to what the binarized model had learned (Fig. S2b), giving evidence for the robustness of the entire framework. Comparing model predictions from the binarized and continuous models showed that the improvement in precision was mostly due to a reranking of false positive variant predictions which were in the proximity of splice sites with intermediate usage levels (Fig. S2c). Stratifying the benchmark to different levels of gene expression showed that the model trained on the continuous usage levels outperformed all other models (including a model trained on gene TPM values instead of splice site usage levels) in all analyzed gene expression regimes (Fig. S3).

Together, these three design improvements – the Pangolin base model, signed features, and continuous splice site usage, on top of the core SpliceMap integration framework – define AbSplice2. Their relative marginal contributions are summarized in Fig. 2c, which shows that the choice of sequence model explains the least variance in performance, while the encoding of tissue-specific splice site usage levels is the dominant driver of prediction improvement. Integrating all these improvements, our final model led to a 2-fold improvement in precision compared to AbSplice1.

The utility of AbSplice lies in tissue-specific predictions. However, in some situations, one may be interested in the overall potential of a variant to cause aberrant splicing in any tissue. While this is not the intended use case of AbSplice, we suggest in this case to consider the maximum effect across all tissues. When benchmarked against tissue-specific splicing outlier prediction in GTEx, the max AbSplice score performs more closely to its underlying sequence-based model Pangolin (Fig. S4). We note however, that variants might still have effects in further contexts not covered by GTEx.

As we have done for AbSplice1, we created AbSplice2-RNA, a model that can further incorporate aberrant splicing events detected from RNA-seq of clinically accessible tissues. This reflects the clinical rare disease diagnostics use case in which RNA-seq could be collected from accessible tissues such as blood or skin fibroblasts, but not from the suspected disease tissue. AbSplice2-RNA further improved the average precision by about 13% (Fig. 2a,b) over the best DNA-only model.

We provide genome-wide predictions for all possible single-nucleotide variants (SNVs) in protein-coding genes for Pangolin and AbSplice2. Additionally, we offer a web interface (https://absplice.cmm.cit.tum.de)^39^ that enables scoring any variant of interest, including insertions and deletions.

### Benchmarking AbSplice2 against recent splicing models

While developing AbSplice2, the general-purpose sequence-to-function model AlphaGenome^24^ was published, which showed strong performance on a variety of tasks including our aberrant splicing benchmark^24^. We therefore compared AbSplice2 to AlphaGenome (AG) as well as one promising other general-purpose model Nucleotide Transformer v3^32^ (NTv3), and SpliceTransformer^33^. Due to computational constraints, we restricted the negative class cases to difficult negatives, i.e. variants with some splice disruptive signals by at least one of SpliceAI, MMSplice, or Pangolin (Methods). This reduced the number of variants from 8 million to 187,733, while not affecting the comparison of the models since only the obvious negative variants were discarded.

AlphaGenome outperformed all other sequence-based models (Fig. S5b,f). Its best individual score (splice-site usage; AP = 0.185) exceeded NTv3’s tissue-specific score (AP = 0.127), SpliceTransformer (AP = 0.084) and AbSplice1 (AP = 0.174), though all remained below AbSplice2 (AP = 0.283).

Given AlphaGenome’s strong performance as a standalone predictor, we asked two related questions: whether SpliceMaps still provide information complementary to AG scores given that AG already predicts tissue-specific splicing remarkably well, and whether AG scores could replace Pangolin as the base model within the AbSplice framework.

To address the first question, we trained two two-feature AbSplice-AG models combining one AG score type with either a binary splice-site expression indicator (as in AbSplice1) or a continuous splice-site coverage feature (as in AbSplice2). Combining the AG score with a binary splice-site expression indicator only marginally improved performance (Fig. S5c,f). This observation is consistent with the AlphaGenome study, where the authors achieved their best performance for aberrant splicing prediction by training a supervised model (effectively an AbSplice model) that combines sequence scores with the binary splice-site expression features introduced in our framework. We found that adding the continuous coverage feature consistently improved performance for all AG score types to a larger extent (Fig. S5c,f). This demonstrates that quantitative information about splice-site coverage remains informative even on top of state-of-the-art tissue-specific predictors, mirroring the advantage of continuous over binary SpliceMap features we observed for SpliceAI and Pangolin.

Turning to the second question, the best AbSplice-AG model (Methods; AP = 0.269) performed slightly worse than AbSplice2 (AP = 0.283; Fig. S5d,f). The current AlphaGenome API does not expose directional gain and loss scores – a distinction that AbSplice2 exploits through Pangolin. When AbSplice2 was trained without directional features, its performance (AP = 0.273) was comparable to the best AbSplice-AG model (Fig. S5d,f), suggesting that this API limitation may account for the remaining gap.

We further asked whether the tissue resolution embedded in the AG scores provides additional benefit once continuous coverage information is included. Combining tissue-agnostic AG scores (maximum absolute value across tissues) with the continuous coverage feature performed equivalently to the corresponding tissue-specific models (Methods; AP = 0.250 vs. 0.252; Fig. S5e,f). AlphaGenome’s ability to generate tissue-specific predictions from sequence alone is a notable property. However, the continuous SpliceMap coverage feature captures this tissue-specific information within the AbSplice framework, making the AG tissue resolution redundant for splicing outlier prediction.

### Improvements of AbSplice2 are confirmed on independent benchmarks

Having established our model in GTEx, we assessed it in another independent cohort consisting of 230 individuals from the BrainGVEx study^40^ with paired whole genome sequence and RNA-seq data from the brain frontal cortex. We applied AbSplice2 using the SpliceMap from GTEx brain frontal cortex (BA9) as a proxy. Aberrant splicing calls from FRASER2 were used to evaluate the predictions. While the model reached similar recall levels as in the corresponding GTEx tissue, the precision of AbSplice in BrainGVEx was reduced compared to GTEx, likely because the reference SpliceMap was not an exact match for the analyzed tissue (SI Fig.6a). Nevertheless, the relative performance of all models was replicated, with AbSplice2 performing best.

In addition to aberrant splicing prediction, we evaluated our model predictions on the task of predicting aberrant gene underexpression in the GTEx dataset. Aberrant splicing often, yet not necessarily, leads to the expression of isoforms containing premature termination codons which can be recognized by the nonsense-mediated decay machinery for degradation. As a result, aberrant splicing predictors are informative about aberrantly underexpressed genes. Therefore, we further evaluated the models at predicting aberrant underexpression, using a benchmark of expression outliers across GTEx that we established earlier^17^. We found that AbSplice2 led to a 2-fold improvement in average precision over AbSplice1. Consequently, replacing AbSplice1 with AbSplice2 in the input features of the aberrant gene underexpression predictor AbExp^17^ led to an improved average precision by about 7% (*P* = 9.3×10^-4^, Fig. 2d).

Building on this observation, we investigated whether AbSplice predictions can aid interpretation of candidate pathogenic variants for rare-disease diagnostics.

To this end, we analysed genome and exome samples from the Solve-RD^41^ cohort, a large cohort spanning a wide range of rare diseases and comprising 24,550 individuals, 15,028 affected and 9,532 unaffected. We defined as rare, all variants with a minor alle frequency less than 0.1% in both the genome and the exome gnomAD databases^42^. We focussed on the SNVs, leveraging our precomputed scores. We first assessed the performance of AbSplice in a tissue-agnostic fashion (i.e., using the AbSplice max score) and analyzing all disorders jointly. We found a mild, yet significant, higher enrichment of high-impact variants in affected individuals for AbSplice2 compared to AbSplice1, SpliceAI, and Pangolin (Fig. S6b). Disorders in the Solve-RD cohort are very heterogeneous and typically not clearly annotated with a primary tissue of interest, at the exception of the neurological or neurodevelopmental disorders (NDD), with 4,780 cases and presumably associated with the central nervous system. Among those cases, we observed a pronounced increase in variant enrichment when restricting the analysis to high-impact variants in NDD-related genes. Notably, the enrichment of high AbSplice2 scores was most significant when using SpliceMaps from the brain and cerebellum (Fig. 2e). These results demonstrate that AbSplice effectively captures clinically-relevant tissue-specific variant effects.

### Dynamically changing splice sites lead to changing predicted variant effects during development

Having established an improved AbSplice model, we next considered extending its application to developmental stages. To this end, we derived SpliceMaps for human RNA-seq data from 7 tissues (brain, cerebellum, heart, kidney, liver, ovary and testis) gathered from 15 different developmental stages spanning embryogenesis to the adult with the earliest recorded stage being 4 weeks post conception^27^. With 1 to 5 samples per tissue-time point pair, the sample size of the developmental dataset was strongly reduced compared to the average 331 samples per tissue in GTEx. Downsampling investigation of the GTEx dataset indicated that SpliceMaps largely agreeing with the original SpliceMaps could still be derived with as few as 4 samples (Fig. S7a) and would not affect AbSplice performance (Fig. S7b). To reach a similar minimal sample size while preserving temporal resolution, we pooled data of each developmental time point and its adjacent time points. Moreover, we confirmed the agreement of the tissue-specific SpliceMaps between the adult stage of the developmental dataset and GTEx by comparing their SpliceMaps (Fig. S7c,d, Jaccard similarity typically above 0.75).

To evaluate how developmental SpliceMaps extend current population-scale resources, we compared detected splice sites in SpliceMaps with the database of SpliceVault (300K-RNA)^30^, which catalogued splice junctions involved with at least one annotated splice sites across 335,663 samples. The AbSplice and SpliceVault frameworks offer complementary clinical utility: AbSplice estimates the overall probability of splicing disruption for a given genetic variant, whereas SpliceVault enables the characterization of the resulting transcriptomic consequences for events anchored to annotated splice sites.

Our comparison revealed that SpliceMaps provide essential coverage for entirely novel junctions – where both the donor and acceptor are unannotated – which are methodologically excluded from the SpliceVault database (Fig. S8). Furthermore, we identified 24,415 splice sites detected across multiple independent individuals and active during organogenesis that remain absent even from high-depth adult population databases (Fig. S8). While SpliceMaps provide high sensitivity by including approximately 1 million rare, stochastic developmental events, the AbSplice model utilizes quantitative usage levels as a continuous predictive feature. This allows the model to naturally distinguish robust signals from stochastic background noise, prioritizing replicated sites while penalizing lower-confidence signals. These findings highlight that while adult population-based catalogs are powerful for characterizing mis-splicing outcomes, developmental SpliceMaps are required to capture the temporal and structural breadth of the human transcriptome.

We computed AbSplice2 scores for all tissues and developmental stages for all possible SNVs within a 100-bp window of all 47,769 previously annotated developmental alternative splicing (devAS) sites^25^. One such devAS site is visualized in Fig. 3a, where exon 6 of the gene *SSBP3* which is known to play a role in embryonic stem cell differentiation and head development^43,44^, is included in an early developmental stage (4 weeks post conception) at a high inclusion level in the brain, but is removed from the spliced transcript at the point of birth and is subsequently missing in the adult brain. In the vicinity of the splice sites of *SSBP3* exon 6 we predicted a significant difference in variant effects between early and late stages of development (Fig. 3a). While there are variants with a high predicted probability to disrupt splicing in early stages, AbSplice2 predicted exclusively low variant impacts in adult stages. One example variant within this region is displayed in Fig. 3b. While the deep learning models Pangolin and MMSplice provided a variant score that was independent of the developmental stage, the splicing metrics from the SpliceMaps captured how the usage of each splice junction changed during development. The drop in inclusion of *SSBP3* exon 6 was captured in the developmental SpliceMaps which was reflected in AbSplice2 predictions being strong in the early stages only. Moreover, the continuous representation of the median splice site usage as opposed to the binary representation used in AbSplice1 led to smoother and more robust predictions over development time (Fig. S9 for an example). This example showcases that while SpliceAI and Pangolin can capture variants affecting splicing, AbSplice2 adds the developmental and tissue context of these effects.

**Fig. 3:**
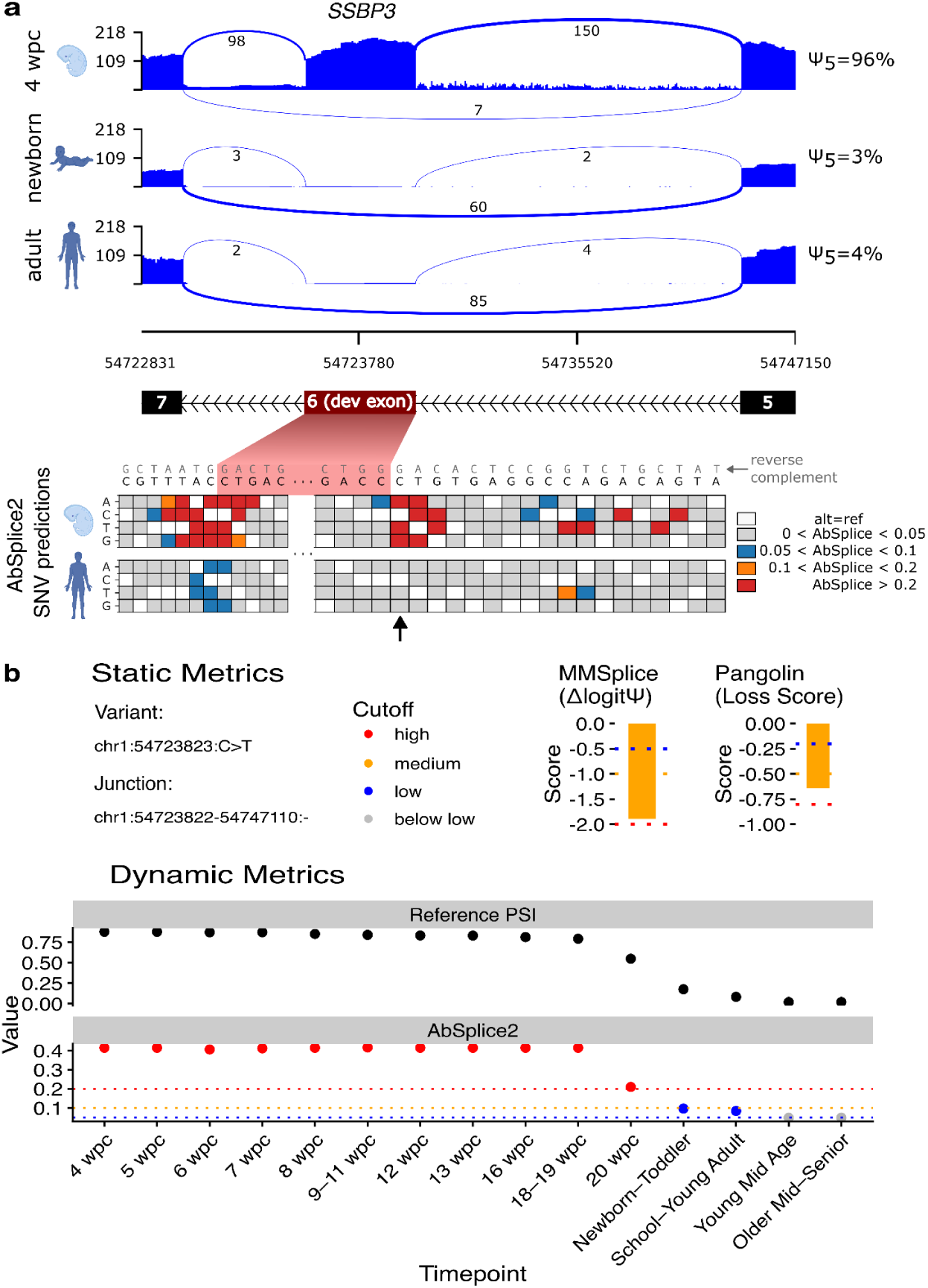
Variants around developmentally dynamic splice sites exhibit developmentally dynamic effects. **a)** Exon 6 in the gene *SSBP3* is expressed in the brain in early stages of development during pregnancy, but not expressed at birth and afterwards. Possible SNVs in the proximity of the donor and acceptor sites of exon 6 lead to high predicted variant effects for early stages of development and low or absent effects for adult stages. 4 wpc: 4 weeks post conception. **b)** Predicted effects of the variant chr1:54723823:C>T (NM_145716.4:c.367-1G>A). The genomic location is indicated in (**a**) with a black arrow (below the heatmap). The deep learning models MMSplice and Pangolin predict a variant effect independent of the developmental timepoint. AbSplice2 predictions change dynamically over time. The drastic drop in the score from early to late developmental stages is caused by the change in the inclusion level of the exon (captured by the SpliceMap metric ‘Reference PSI’). Timepoints are labelled according to the developmental study by Mazin et al.^25^. Scores are colored based on applied cutoffs: high (AbSplice2: 0.2, Pangolin: 0.8, MMSplice: 2), medium (AbSplice2: 0.1, Pangolin: 0.5, MMSplice: 1), low (AbSplice2: 0.05, Pangolin: 0.2, MMSplice: 0.5).

We predicted for each tissue the number of variants genome-wide that led to a splice-disrupting effect in an early stage of development (from conception to toddler age) and no effect in the adult (Fig. 4a). The brain, which is the tissue with most devAS events^25^, showed the highest number of such variants (N=18,206 which is 10% of all high impact variants in the brain across all developmental stages around devAS sites, Fig. 4a). These variants may be of clinical importance as developmentally changing splicing events were shown to be highly conserved across evolution^25^. Fitting this hypothesis, these development-specific effect variants are enriched in loss-of-function (LoF) intolerant genes, with genes in the 10% most LoF-intolerant group being 6.8-fold more likely to harbor a development-specific effect variant compared to the 10% most LoF-tolerant genes (Fisher’s exact test *P* = 6.7×10^-65^) (Fig. 4b). Moreover, these variants were also significantly enriched in NDD genes^34^ (Fig. 4c).

**Fig. 4:**
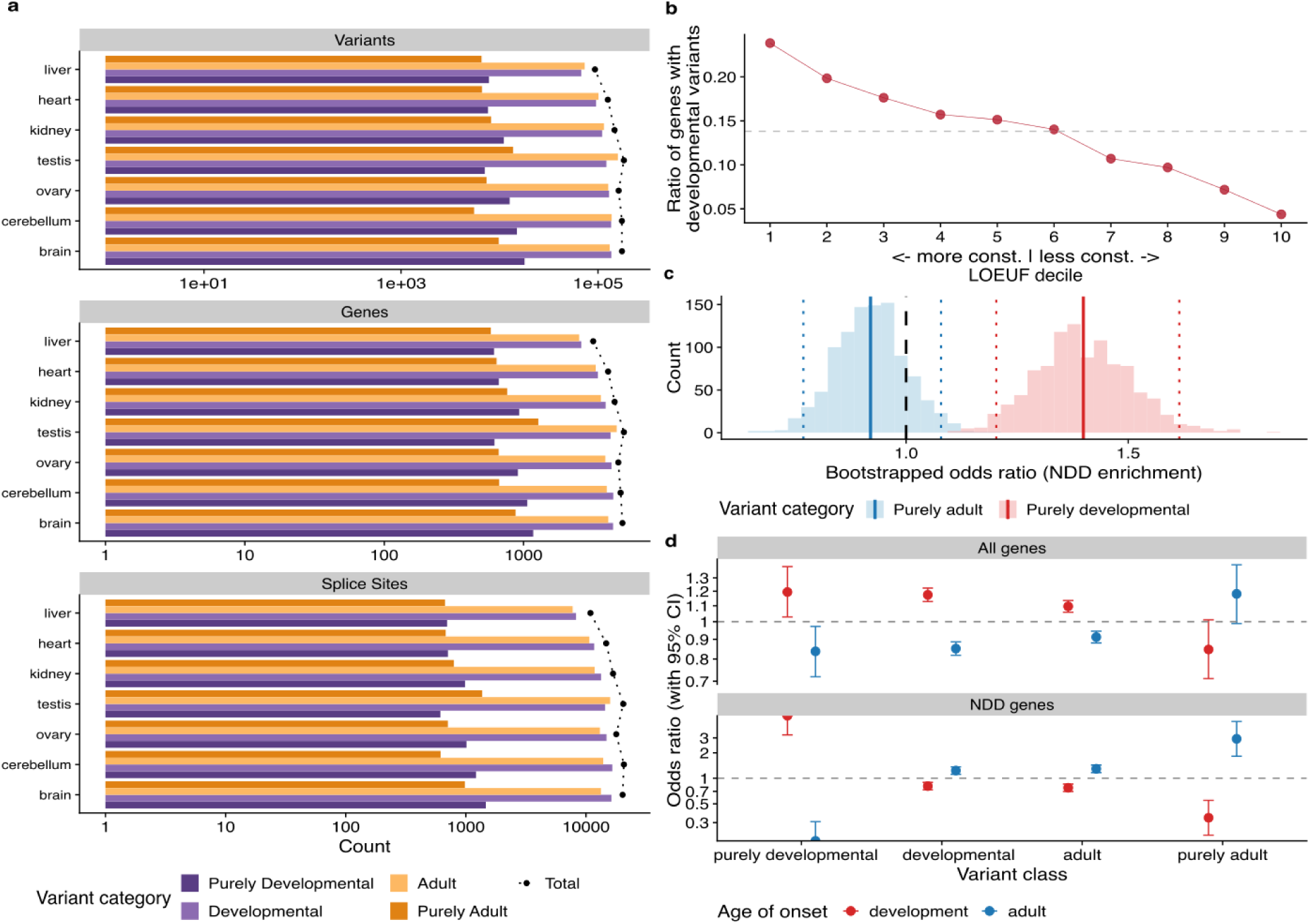
Purely developmental variants are enriched in functionally relevant genes and individuals with an early age of disease onset. **a)** All possible SNVs around devAS sites were scored with AbSplice2 for 15 developmental stages (including early development and adult stages) and 7 tissues. Variant categories were defined based on AbSplice2 scores. The category ‘Total’ contains all variants with a score above 0.2 in at least one of the 15 developmental stages. The category ‘Developmental’ contains all variants with a score above 0.2 in at least one early developmental stage (from embryogenesis to toddler). The category ‘Purely Developmental’ contains all variants with a score above 0.2 in at least one early developmental stage (from embryogenesis to toddler) and a score below 0.05 for adult stages. The category ‘Adult’ contains all variants with a score above 0.2 in at least one adult stage. The category ‘Purely Adult’ contains all variants with a score above 0.2 in at least one adult stage and a score below 0.05 for all early developmental stages (from embryogenesis to toddler). Top: The number of variants per tissue for the different variant categories. Middle: The number of unique genes affected by those variants. Bottom: The number of unique splice sites affected by those variants. **b)** Ratio of genes containing a developmental variant (variants with AbSplice2 above the high cutoff (0.2) at any developmental stage pre-birth up to toddler and AbSplice2 below the low cutoff (0.05) for adult stages) as a function of LOEUF decile. Genes in lower LOEUF deciles are more constrained. The dotted grey line indicates the genome-wide baseline ratio of genes with developmental variants. **c)** Enrichment of purely developmental and purely adult variants in genes related to neurodevelopmental disorders. Displayed is the histogram of odds ratios based on 1,000 bootstrapped gene set samples. The solid lines represent the mean of the distributions. The dotted lines represent the 95% confidence intervals. **d)** Enrichment of purely developmental, developmental, adult and purely adult variants in individuals within Solve-RD affected with neurological or neurodevelopmental disorders. Brain-specific scores (maximum over brain and cerebellum) from AbSplice2 are used. Odds ratios were computed for the different variant classes and individuals with early (from embryogenesis to toddler) and adult (above 40 years) age of onset for all genes (top) and neurodevelopmental disorder (NDD) genes (bottom). Confidence intervals are based on fisher tests.

Analyzing individuals affected with neurological and neurodevelopmental disorders in the Solve-RD cohort, we observed that purely developmental high-impact variants were enriched in individuals with an early age of onset, while in contrast, purely adult variants showed an enrichment in individuals with an age of onset during adulthood (Fig. 4d). This correlation with age of onset was further strengthened when restricting the analysis to NDD-related genes (Fig. 4d). This demonstrates that the temporal dynamics predicted by AbSplice provide clinically relevant context for unravelling the developmental context of disease onset.

### Application to rare disease samples

To assess the clinical potential of predicting developmental aberrant splicing, we applied AbSplice2 to 64,736 rare disease participants from the UK National Genomics Research Library^45^ (NGRL). We focussed on variants located within 100 bp of devAS splice sites and for which AbSplice2 predicted an effect during early development (AbSplice2 score above the cutoff 0.05) that is stronger than in adult. We further restricted to variants in ‘Green’ genes confidently linked with genetic disorders from PanelApp^46^, identifying a total of 26 unique variants in 47 individuals (Table 1, SI Table 1). One example is a splice acceptor variant in *ARFGEF1* (chr8:67203252:C>T, NM_006421.5:c.4960-1G>A) that had already been identified as the causal variant in an individual recruited with intellectual disability. This gene is associated with developmental delay, epilepsy, and intellectual disability. AbSplice2 predicted this variant to activate a weak splice site within the adjacent exon causing a frameshift with an effect that is strongest in the brain during an early neurodevelopmental window (between 12 and 16 weeks post-conception), to persist at a high level into young adulthood and to be absent in older adults (Fig. S10a). ADP ribosylation factor guanine nucleotide exchange factor (*ARFGEF*) genes are involved in cellular trafficking and neuronal development in the hippocampus, and play an important role in neurite development and axon elongation^47^ with evidence suggesting that neurite development spans from early development into adolescence, with the possibility of different tissues having higher or lower neurite densities at different timepoints^48^.

Three rare (gnomAD allele frequency < 10^-4^) variants in three individuals were prioritised as potential diagnostic candidates after reviewing phenotypic fit and mode of inheritance (Table 1). The first variant (chr8:38428099:C>T, NM_023110.3:c.449-6G>A) lies in *FGFR1* in an individual with hypogonadotrophic hypogonadism. This variant introduces an AG dinucleotide in the AG exclusion zone upstream of the canonical acceptor site^49^, causing a frameshift and introduction of a premature termination codon (Fig. 5a). This variant is classified in ClinVar as having ‘conflicting interpretations of pathogenicity’ with both likely pathogenic, and uncertain significance classifications. *FGFR1* plays a crucial role in the development of the neuroendocrine hypothalamus and is central to the genesis of gonadotropin-releasing hormone (GnRH) secreting neurons in the olfactory bulb, and their migration into the hypothalamus during development^50^. Variants that result in a loss of function in the *FGFR1* gene have been found to cause a failure of neuronal migration and reduced GnRH secretion^50^ leading to hypogonadotropic hypogonadism and anosmia (Kallmann syndrome)^51^. AbSplice2 predicted embryogenesis-specific activation of this acceptor site in early forebrain development (Fig. 5b) and in the kidney, but not after birth. The effect in the forebrain is consistent with the disease mechanism.

**Fig 5:**
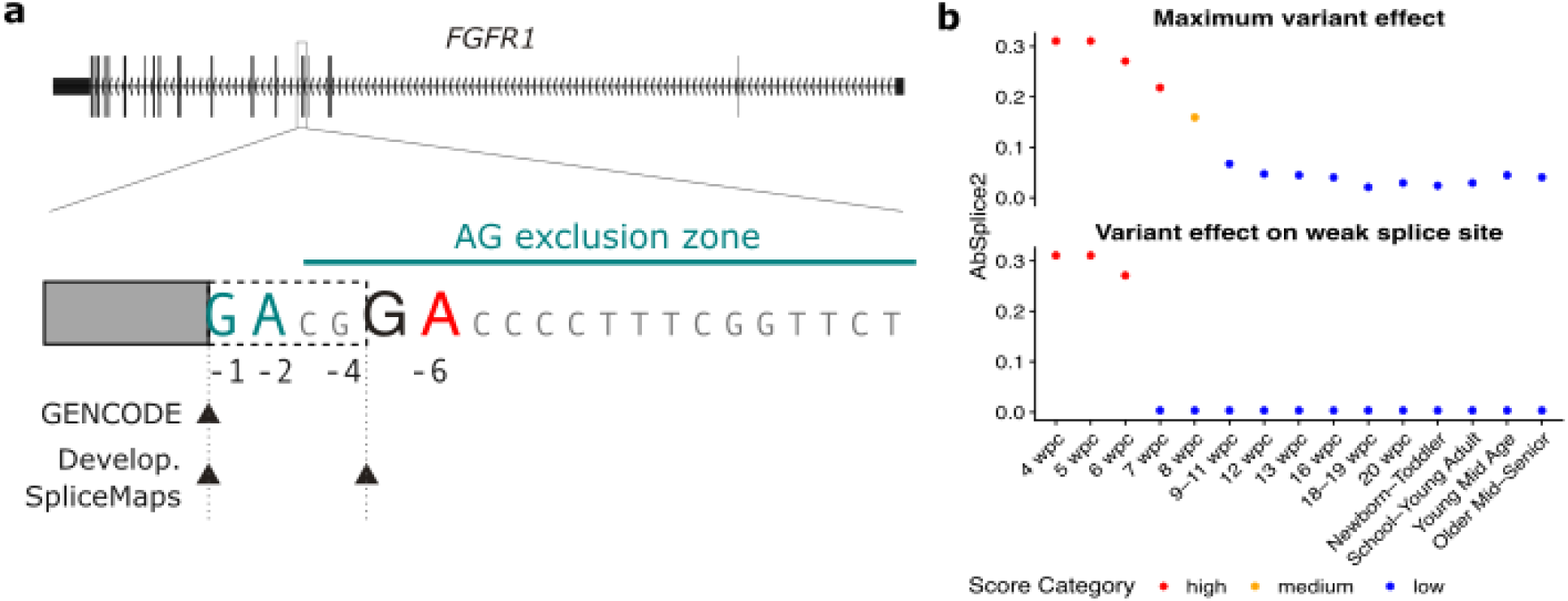
AbSplice2 predicts the activation of a weak splice site in early development providing a potential diagnostic candidate variant for an undiagnosed individual. **a)** Acceptor-site creating variant (chr8:38428099:C>T, NM_023110.3:c.449-6G>A) in the AG exclusion zone of exon 6 of the *FGFR1* gene that was identified in an individual within the NGRL recruited with hypogonadotrophic hypogonadism. Variants that result in a loss of function in *FGFR1* have been found to lead to the disease phenotype (Kallmann syndrome). The variant affects a weak splice site identified during embryogenesis (Develop. SpliceMaps, lower track) but not in the GENCODE annotation, which would result in a frameshift. **b)** Top. Maximum AbSplice2 effect predictions of the variant regarding loss of the canonical site and the gain of the weak splice site. Bottom. AbSplice2 predictions of the variant on the weak splice site. The effect is predicted to be strong only in the weeks 4-6 post conception.

The second candidate variant was identified in *FN1*, a gene associated with skeletal dysplasia. Whilst we identified an elevation in AbSplice2 scores at early developmental stages across tissues, the specific tissues with the greatest relevance to the disorder (skeletal dysplasia) were unavailable within our data (Fig. S10b). For *NDUFS2*, a mitochondrial complex 1 subunit associated with Leigh syndrome, the clinical relevance in the NGRL participant remains uncertain, as deleterious variants in this gene are typically associated with early-onset fatal outcomes in childhood, while this participant is an adult. It is possible that this variant may be a hypomorphic non-coding variant, and therefore result in a less severe phenotype, however we are unable to confirm this currently. All remaining variants did not represent good candidates for the individual’s disorder when mode of inheritance or phenotypic matching were considered, including 8 instances where the participants’ case had been solved, or partially solved with the causal variant being found in a different gene (Table S1).

## Discussion

Here, we have developed a model for tissue-specific aberrant splicing prediction across developmental stages. Thereby, our AbSplice framework predicts a class of variants whose effect on splicing changes over the course of development. Additionally, we improved on the previous AbSplice model by replacing SpliceAI with Pangolin, leveraging the directionality of predicted splicing changes as well as the predicted affected position, which led to improved precision. Second, we transitioned from a binary quantification of splice site usage to continuous usage levels which provided more robust and precise predictions. Moreover, we showed how AbSplice aids the interpretation of rare variants for rare disease diagnostics by indicating for which developmental stages and tissues a variant can affect splicing. We provide the code and developmental SpliceMaps spanning stages from embryogenesis to adulthood. For convenience, we also provide precomputed scores for AbSplice2 and Pangolin for all possible SNVs genome-wide.

In our previous publication^14^ we used FRASER aberrant splicing calls as ground truth to train and evaluate AbSplice, as we found FRASER to be more robust and precise than other outlier callers^9,10^. In this study we employed an improved ground truth based on FRASER2, which reduced initial outlier calls by nearly six-fold. We found that model performance was robust to this change in ground truth, primarily because our rare-variant-based filtering had already suppressed most noise from the original outlier caller. Retraining on FRASER2 yielded only marginal performance differences and did not alter the learned feature contribution functions of the AbSplice model. These results demonstrate that the model captures a consistent biological signal across different ground truth sources, validating the reliability of the AbSplice framework and the previously published AbSplice model.

The developmental predictions were based on SpliceMaps constructed from limited sample sizes and only for seven tissues. While we showed that the available sample sizes led to reasonably robust estimates, there are potential benefits of larger cohorts for more robust and comprehensive splicing quantifications. We anticipate that efforts such as the upcoming developmental data from GTEx will enable the construction of more reliable SpliceMaps and facilitate the detection of weakly spliced junctions, which are often present in only a few individuals, but can lead to drastic aberrant splicing events when activated by a variant^30,52^. Moreover, an update would only require computing SpliceMaps from such data owing to the modular structure of AbSplice. Our evaluations in this study and our previous study^14^ showed that the AbSplice framework is very robust to changes in the SpliceMaps while keeping the model unchanged.

We applied AbSplice to rare disease participants from the UK National Genomics Research Library (NGRL) and prioritized variants predicted to have a developmental splicing effect, resulting in 26 candidate variants in known disease-associated genes. While AbSplice predicted developmental splicing disruptions, not all identified variants were in genes clearly related to the individual’s specific phenotypes. Many variants were heterozygous in genes that require a biallelic impact to cause disease. While these were not considered to be causal for the individual’s phenotypes, this does not imply that the underlying splicing predictions were incorrect. It remains possible that the predicted splicing alterations are real, but that disrupted splicing is not the primary mechanism underlying the observed phenotype – or that it contributes without being sufficient on its own. For some variants we observed evidence of weak splicing at the affected site exclusively in the developmental dataset, but not in adult GTEx tissues, demonstrating the importance of considering temporally dynamic regulatory patterns in variant interpretation. These findings underscore the added value of incorporating developmental context into variant interpretation.

AbSplice helps narrow down splicing-relevant variants to specific tissues and developmental stages for which SpliceMaps are available. Consequently, using the maximum score across all tissues is not the intended usage and leads to reduced precision in tissue-specific tasks (Fig. S4). Moreover, since not all tissues or developmental stages are covered, variants might still have effects in unobserved contexts. Therefore, we see AbSplice as a companion tool to link predicted variant effects to the observed phenotypes by analyzing the developmental and tissue-specific prediction^53–55^.

One limitation of our approach is that it depends on the sensitivity of the underlying sequence-based predictor. If the model fails to predict a splicing event, our framework cannot recover the prediction even in regions where a tissue-specific splice site is annotated. This could be particularly problematic for tissue-specific or developmental splicing events underrepresented in the training data of sequence-based models. Nevertheless, recent saturation mutagenesis studies have shown that sequence-based models such as SpliceAI and Pangolin can accurately predict the effects of mutations in splicing regulatory elements and recapitulate regulatory architectures of alternative exons^56,57^. These findings support the use of sequence-based models to estimate the effects of cis-regulatory variants, which can subsequently be contextualized using empirical SpliceMaps, as in AbSplice. However, unlike end-to-end sequence-based predictors, this approach does not capture the genetic determinants of context-specificity of splicing. In agreement with the authors of Alphagenome, our evaluation shows that AlphaGenome makes a significant step forward in capturing tissue-specific splicing, yet SpliceMaps remain helpful. Future progress will likely require both, new modeling techniques, for instance building on self-supervised^58–62^ or supervised^24,63–65^ foundation models, along with expanded training datasets covering more tissues, cell types, and developmental stages.

## Data availability

No new data was generated for this study. SpliceMaps for all 49 GTEx tissues (hg19 and hg38) and all 15 developmental timepoints (hg19) in brain, cerebellum, heart, kidney, liver, ovary and testis are available at Zenodo; hg19 SpliceMaps: https://doi.org/10.5281/zenodo.7821508, hg38 SpliceMaps: https://doi.org/10.5281/zenodo.6387937. Precomputed Pangolin scores (hg38) are available at Zenodo, https://doi.org/10.5281/zenodo.15649337. Precomputed AbSplice2 scores (hg38) in all 49 GTEx tissues are available at Zenodo,https://doi.org/10.5281/zenodo.15644547. Precomputed AbSplice2 scores within 100 bp of devAS sites for all 15 developmental timepoints (hg19) are available at Zenodo, https://doi.org/10.5281/zenodo.15674611. The Solve-RD dataset is available in EGA (study EGAS00001003851) and can be accessed following approval from the Solve-RD Data Access Committee. Due to potential donor re-identification when revealing rare variants, the GTEx benchmark dataset cannot be shared without restrictions. Users with access to the GTEx data can reproduce the benchmark using the code repository below. The GTEx v8 dataset is available at (under dbGaP protection) https://gtexportal.org/home. The BrainGVEx dataset is available at the PsychENCODE portal (SynID: syn459090949). Research on the de-identified patient data used in this publication can be carried out in the Genomics England Research Environment subject to a collaborative agreement that adheres to patient led governance. All interested readers will be able to access the data in the same manner that the authors accessed the data. For more information about accessing the data, interested readers may contact or access the relevant information on the Genomics England website: https://www.genomicsengland.co.uk/research.

## Code availability

AbSplice2 predictions using the tissue-specific SpliceMap annotations can be performed with the custom-written python package ‘absplice’ (publicly available at: https://github.com/gagneurlab/absplice2). The analyses are available at: https://github.com/gagneurlab/AbSplice2_analysis.

## Declaration of generative AI and AI-assisted technologies in the manuscript preparation process

During the preparation of this work the authors used Gemini (Google) in order to refine the phrasing and clarity of the manuscript, and Claude (Anthropic) for coding assistance. After using these tools, the authors reviewed and edited the content as needed and take full responsibility for the content of the published article.

## Acknowledgements

We thank all the scientists who isolated and sequenced the data which builds the basis for our research. We thank the authors of Pangolin for making their code easily available and the authors of the developmental splicing study for providing such complete and well documented Supplementary data. We thank Alex Karollus, Pedro Tomaz da Silva, Johannes Hingerl, Laura Martens and Muhammed H. Çelik for insightful discussions and advice. We thank Christian Mertes for his insights on implementing the web interface. The Ethics Committee of the Technical University of Munich issued a statement of no objection regarding this study (reference #2025-263-S-KK). This study was funded by the Deutsche Forschungsgemeinschaft (DFG, German Research Foundation) – Project-ID 461264291. This study was funded by the Deutsche Forschungsgemeinschaft (DFG, German Research Foundation) via the project NFDI 1/1 “GHGA - German Human Genome-Phenome Archive” (#441914366 to N.Wagner, AMN, VAY, JG), the German Bundesministerium für Bildung und Forschung (BMBF) through the ERA PerMed project PerMiM (01KU2016B to N.Wagner, VAY, JG), the Helmholtz Association under the joint research school Munich School for Data Science - MUDS (to N.Wagner and SL). This study was funded by the European Research Council (ERC) (EPIC, Grant number: 101118521) funded by the European Union. ERDERA has received funding from the European Union’s Horizon Europe research and innovation programme under grant agreement N°101156595. Views and opinions expressed are those of the author(s) only and do not necessarily reflect those of the European Union or any other granting authority, who cannot be held responsible for them. Parts of Fig. 1 and Fig. 3 were created with BioRender.com. The Genotype-Tissue Expression (GTEx) project was supported by the Common Fund of the Office of the Director of the National Institutes of Health, and by the NCI, NHGRI, NHLBI, NIDA, NIMH and NINDS. The funders (of GTEx) had no role in study design, data collection and analysis, decision to publish or preparation of the manuscript. N.Whiffin is supported by a Sir Henry Dale Fellowship jointly funded by the Wellcome Trust and the Royal Society (grant no. 220134/Z/20/Z) and a Lister Institute Research Prize. This study makes use of data collated and/or generated by the Solve-RD project, derived from the datasets EGAD00001009767, EGAD00001009768, EGAD00001009769 and EGAD00001009770. The Solve-RD project has received funding from the European Union’s Horizon 2020 research and innovation programme under grant agreement No 779257. This study was supported by the European Reference Networks ERN [EURO-NMD, RND, ITHACA, RITA, EpiCARE, GENTURIS] (https://ec.europa.eu/health/ern/networks_en).

This research was made possible through access to data in the National Genomic Research Library, which is managed by Genomics England Limited (a wholly owned company of the Department of Health and Social Care). The National Genomic Research Library holds data provided by patients and collected by the NHS as part of their care and data collected as part of their participation in research. The National Genomic Research Library is funded by the National Institute for Health Research and NHS England. The Wellcome Trust, Cancer Research UK and the Medical Research Council have also funded research infrastructure.

## Author information

### Contributions

N.Wagner and J.G. conceived the study. N.Wagner and A.M.N. designed the model and implemented the framework. N.Wagner, A.M.N., A.M.G., S.L., C.S. and V.A.Y. analyzed the data. A.M.G. performed the Genomics England NGRL analysis. A.M.N. implemented the web interface. N.Wagner and J.G. wrote the manuscript with input from all the authors. J.G. and N.Whiffin supervised the project.

