## Supplementary Material for "Aberrant splicing prediction during human organ development"

| Variant | Participant Count | Gene | Mode of Inheritance | gnomAD AC/AN/AF | Participant recruited disorder(s) | Gene associated disorder | Notes | AbSplice2 observations |
|---|---|---|---|---|---|---|---|---|
| chr1:181793663A>C (NM_001205293.3:c.5899-2A>C, hg19: chr1:181762799A>C) | 1 | *CACNA1E* | Autosomal Dominant | 2/1608198/1.24E-06 | Paediatric disorders | *CACNA1E*-related epileptic encephalopathy with contractures, macrocephaly, and dyskinesia (G2P) | Heterozygous. Case solved CHD7 | Strong dev effects in brain and cerebellum at junction chr1:181759692-181762800:+ |
| chr2:99554536T>C (NM_001386135.1:c.3336-2A>G, hg19: chr2:100170998T>C) | 1 | *AFF3* | Autosomal Dominant | 22/1611672/1.37E-05 | Coarse facial features including Coffin-Siris-like disorders | *AFF3*-related KINSSHIP syndrome, *AFF3*-related intellectual disability (G2P) | Heterozygous. Case solved AUTS2 | Strong dev effects in brain and kidney at junction chr2:100170996:100171144:- |
| chr5:179623635C>G (NM_001257293.2:c.-72G>C, hg19: chr5:179050636C>G) | 1 | *HNRNPH1* | Autosomal Dominant | Not in gnomAD | Intellectual disability | *HNRNPH1*-related neurodevelopmental disorder (G2P) | Heterozygous. MYT1L VUS is better match | Strong dev effects in brain and testis at junction chr5:179044933-179050646:- |
| chr11:57806454C>G (NM_001085458.2:c.1877-7C>G, hg19: chr11:57573926C>G) | 1 | *CTNND1* | Autosomal Dominant | 7/1601964/4.37E-06 | Intellectual disability | *CTNND1*-related blepharo-cheiro-dontic syndrome (G2P) | Heterozygous. Unclear phenotype match | Strong dev effects in brain, cerebellum, heart, kidney, ovary and testis at multiple junctions. Affected junctions: chr11:57573507:57573932:+ (brain, cerebellum, heart, kidney, ovary, testis), chr11:57572252:57573932:+ (brain), chr11:57559145:57573932:+ (brain), chr11:57569668-57573932:+ (cerebellum), chr11:57509426-57573932:+ (cerebellum) |
| chr12:57582586A>G (NM_004984.4:c.2993-16A>G, hg19: chr12:57976369A>G) | 1 | *KIF5A* | Autosomal Dominant | 5/1608500/3.11E-06 | Noonan syndrome | *KIF5A*-related severe neonatal myoclonus (G2P) | Heterozygous. Case partially solved USP7 variant is better match | Strong dev effects in brain and cerebellum at multiple junctions |
| chr15:57252494T>A (NM_207037.2:c.1260+2T>A, hg19: chr15:57544692T>A) | 1 | *TCF12* | Autosomal Dominant | 7/1612868/4.34E-06 | Paediatric disorders | *TCF12*-related neurodevelopmental disorder with coronal craniosynostosis (G2P) | Heterozygous. Case solved HNRNPU | Strong dev effects in all tissues at junction chr15:57544690:57545459:+ |
| chr17:31253005G>A (NM_001042492.3:c.4173+5G>A, hg19: chr17:29580023G>A) | 1 | *NF1* | Autosomal Dominant | 1/1610582/6.21E-07 | Paediatric disorders | *NF1*-related neurofibromatosis | Heterozygous, Actionable variant identified in CACNA1F is better match | Strong dev effects in brain at junction chr17:29580018-29585361:+ and chr17:29580018:29685497:+ |
| chr22:23766253T>G (NM_213720.3:c.284A>C, hg19: chr22:24108440T>G) | 2 | *CHCHD10* | Autosomal Dominant | 8894/988434/9.00E-03 | IUGR and IGF abnormalities, Early onset dystonia | Frontotemporal dementia and/or amyotrophic lateral sclerosis 2, Mitochondrial disease (ClinGen) | Heterozygous, ClinVar Likely Benign | Strong dev effects in heart at junction chr22:24108429-24109560:- |
| chr15:89303919A>G (NM_001113378.2:c.3058+4A>G, hg19: chr15:89847150A>G) | 1 | *FANCI* | Autosomal Recessive | 39/1613850/2.42E-05 | Intellectual disability | *FANCI*-related Fanconi anemia (G2P) | Pathogenic variant found in LZTR1 is better match | Strong dev effects in multiple junctions in brain and cerebellum |

| | | | | | | | | |
|---|---|---|---|---|---|---|---|---|
| chr16:75542598T>C (NM_001077418.3:c.664+4A>G, hg19: chr16:75576496T>C) | 3 | *TMEM231* | Autosomal Recessive | 78/1613500/4.83E-05 | Retinal disorders, Cerebellar hypoplasia, Rod-cone dystrophy | *TMEM231*-related Joubert syndrome (G2P) | Heterozygous, no biallelic candidate variants identified. | Strong dev effects in brain and cerebellum at junction chr16:75575353:75576499:- |
| chr16:75542601C>T (NM_001077418.3:c.664+1G>A, hg19: chr16:75576499C>T) | 1 | *TMEM231* | Autosomal Recessive | 4/1613706/2.48E-06 | Cystic renal disease | *TMEM231*-related Joubert syndrome (G2P) | Heterozygous, no biallelic candidate variants identified. | Strong dev effects in brain and cerebellum at junction chr16:75575353:75576499:- |
| chr9:13147659C>G (NM_001378778.1:c.3631-1G>C, hg19: chr9:13147658C>G) | 1 | *MPDZ* | Autosomal Recessive | 21/1607054/1.31E-05 | Hereditary ataxia | *MPDZ*-related nonsyndromic hydrocephalus (G2P) | Heterozygous, no biallelic candidate variants identified. | Strong dev effects in brain, heart, kidney, ovary and cerebellum at junction chr9:13147657-13150509:- |
| chr9:89334732T>C (NM_024077.5:c.1089+2T>C, hg19: chr9:91949647T>C) | 1 | *SECISBP2* | Autosomal Recessive | 1/1608240/6.22E-07 | Paediatric disorders | *SECISBP2*-related thyroid hormone metabolism, abnormal (G2P) | Heterozygous, no biallelic candidate variants identified. | Strong dev effects in brain at junction chr9:91949564:91953372:+ |
| chr10:60166590C>A (NM_020987.5:c.2614+1G>T, hg19: chr10:61926348C>A) | 1 | *ANK3* | Autosomal Recessive | 11/1612514/6.82E-06 | Intellectual disability | Intellectual disability (ClinGen) | Heterozygous. Does not segregate in family | Strong dev effects in heart at junction chr10:61898845-61926348:- |
| chr7:108237751C>T (NM_001037132.4:c.124+1G>A, hg19: chr7:107878195C>T) | 1 | *NRCAM* | Autosomal Recessive | 21/1592558/1.32E-05 | Paediatric disorders | *NRCAM*-related neurodevelopmental disorder with dysmorphic features, hypotonia, and spasticity (G2P) | Heterozygous, no biallelic candidate variants identified. | Strong dev effects in brain and cerebellum at junction chr7:107875132-107878195:-, |
| chr8:144085277T>G (NM_003801.4:c.1261-12T>G, hg19: chr8:145140180T>G) | 2 | *GPAA1* | Autosomal Recessive | 37/1588380/2.329e-5 | Hereditary ataxia, Ultra-rare undescribed monogenic disorders | *GPAA1*-related developmental delay, epilepsy, cerebellar atrophy, and osteopenia (G2P) | Heterozygous. ClinVar Likely Benign | Strong dev effects in heart and liver at junction chr8:145140041-145140191:+, Some dev effects in brain at junction chr8:145139512-145140191:+ |
| chr1:46190809T>G (NM_017739.4:c.1540-25A>C, hg19: chr1:46656481T>G) | 1 | *POMGNT1* | Autosomal Recessive | 22/1584768/1.39E-05 | Congenital myopathy | *POMGNT1*-related muscular dystrophy-dystroglycanopathy congenital with brain and eye anomalies/ intellectual developmental disorder, type B, 3/ limb-girdle (G2P) | Heterozygous, no biallelic candidate variants identified. | Strong dev effects in multiple junctions in all tissues |
| chr2:206149017T>C (NM_005006.7:c.338+3A>G, hg19: chr2:207013741T>C) | 1 | *NDUFS1* | Autosomal Recessive | 42/1601574/2.62E-05 | Epilepsy plus other features | *NDUFS1*-related Leigh syndrome, *NDUFS1*-related mitochondrial respiratory chain complex I deficiency (G2P) | Heterozygous, no biallelic candidate variants identified. | Strong dev effects in brain at junction chr2:207012558:207013743:- |
| chr4:127891683T>C (NM_014264.5:c.2038+2T>C, hg19: chr4:128812838T>C) | 1 | *PLK4* | Autosomal Recessive | 0/1276152/0.00E+00 | Paediatric disorders | *PLK4*-related microcephaly, growth failure and retinopathy (G2P) | Heterozygous. Case solved SOS1 | Strong dev effects in brain, cerebellum, ovary, heart and liver at junction chr4:128812836-128813519:+. In testis opposite dev effects. |
| chr5:6606872C>T (NM_017755.6:c.1549G>A, hg19: chr5:6606985C>T) | 1 | *NSUN2* | Autosomal Recessive | Not in gnomAD | Intellectual disability | *NSUN2*-related intellectual disability (G2P) | Heterozygous. Poor match with specific phenotype | Dev effects in brain and cerebellum at multiple junctions |

| | | | | | | | | |
|---|---|---|---|---|---|---|---|---|
| chr3:51987627A>T (NM_000666.3:c.921+3A>T, hg19: chr3:52021643A>T) | 1 | *ACY1* | Autosomal Recessive | 1/1613852/6.20E-07 | Intellectual disability | *ACY1*-related aminoacylase 1 deficiency (G2P) | Heterozygous, no biallelic candidate variants identified. | Minor dev effects |
| chr1:21838825C>T (NM_005529.7:c.10150G>A, hg19: chr1:22165318C>T) | 1 | *HSPG2* | Autosomal Recessive | 31/1611812/1.92E-05 | Likely inborn error of metabolism | *HSPG2*-related Schwartz-Jampel syndrome, *HSPG2*-related dyssegmental dysplasia, Silverman-Handmaker type (G2P) | Heterozygous. Not good phenotypic match | Minor dev effects |

**Table S1:** 22 candidate variants identified within the UK National Genomics Research Library (NGRL), including the number of individuals in the NGRL observed with that variant (GEL Count); the allele frequency in gnomAD v.4, the mode of inheritance and gene-associated disorder as recorded in DECIPHER, the disorder for which the participant was recruited, the disorder most closely linked to the gene as provided by DECIPHER from the Gene2Phenotype (G2P) and ClinGen databases, notes on prioritisation including existing identified alternate candidate genes, and how well HPO terms recorded for the AbSplice2 variant gene match with the individual's phenotype, as well as information on AbSplice2 predictions across tissues and developmental stages. Where an existing variant has been identified, AbSplice variants were assessed with reference to whether they could further contribute to an explanation of the individual's phenotype. Junction coordinates in AbSplice2 observations are in hg19.

### Supplemental Figures

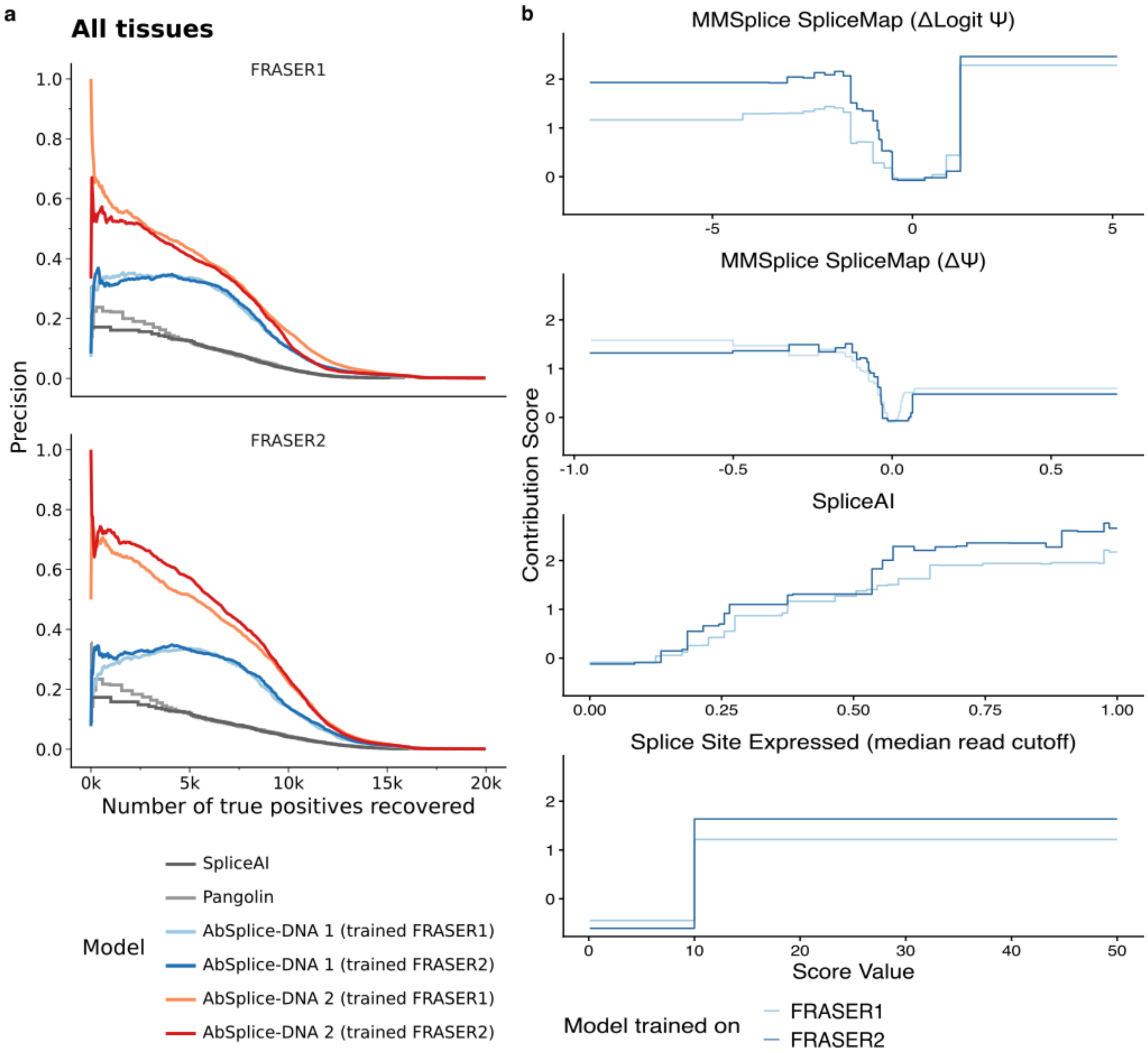


**Figure S1: AbSplice is robust to the splicing outlier caller used for training.**
**a)** Precision-recall curves for models evaluated against FRASER1 (top) and FRASER2 (bottom) as ground truth, aggregated across all GTEx tissues (n = 49). Models include SpliceAI, Pangolin, AbSplice-DNA 1, and AbSplice-DNA 2, each trained on either FRASER1 or FRASER2 ground truth. **b)** Feature contribution functions (the higher the more important to the overall probability prediction[63]) of AbSplice1 models trained on FRASER1 and FRASER2 ground truth. The feature contributions did not change strongly while changing the ground truth.

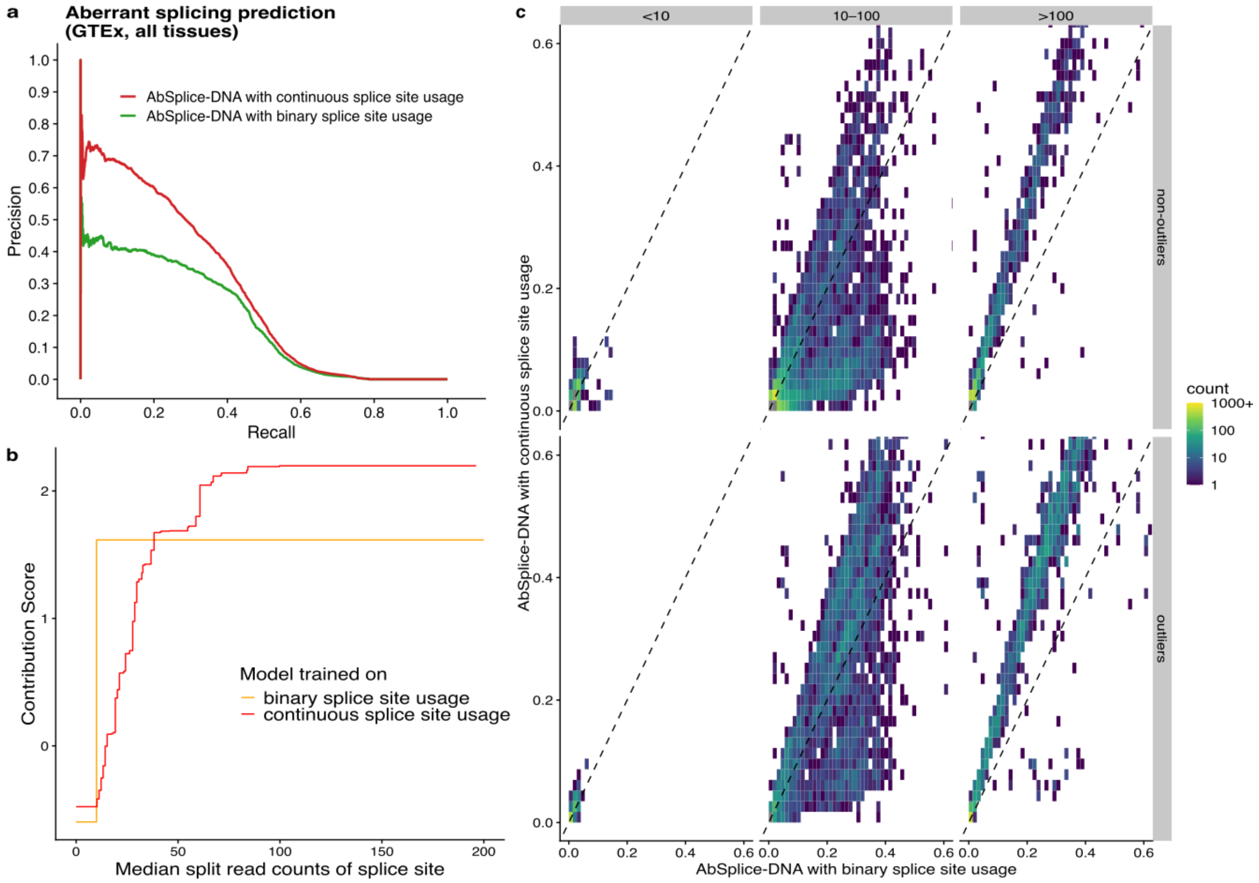


**Figure S2: Using continuous splice site usage improves model performance. a)** Precision–recall curve comparing the overall prediction performance on all GTEx tissues of AbSplice models trained on a binary feature from SpliceMap indicating if the splice site is expressed in the target tissue (using a cutoff of 10 reads for the median number of split reads sharing the splice site) as well as on a continuous feature (the median number of split reads sharing the splice site). The other input features of the two models are identical and include the signed variant effect scores of MMSplice and Pangolin. **b)** Comparison of the learned feature contribution functions of AbSplice models shown in (**a**) for the binary and continuous splice site usage feature. **c)** Comparison of AbSplice model scores shown in (**a**) for outliers (bottom row) and non-outliers (top row) in different splice site usage regimes (columns, median split read counts with the two cutoffs 10 and 100 match the break points in the binary splice site usage function depicted in (**b**)). Non-outliers were randomly subsampled to match 100 times the number of outliers for visualization purposes.

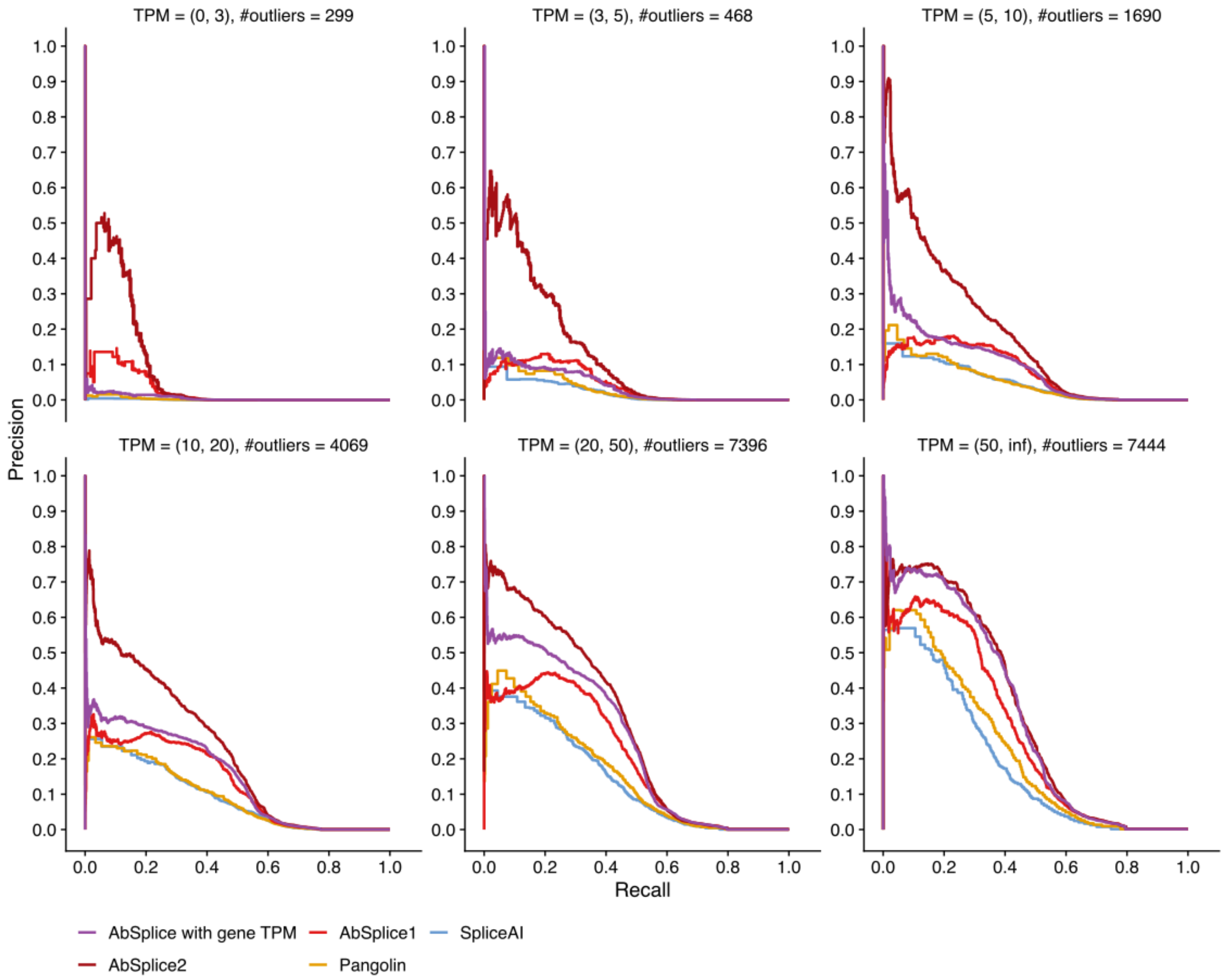


**Figure S3: AbSplice model trained on continuous splice site usage levels outperforms other models in all gene expression regimes.** Precision–recall curves comparing the overall prediction performance on all GTEx tissues of different models constrained for different gene expression regimes. Displayed are SpliceAI, Pangolin, AbSplice1, AbSplice2 and AbSplice2 with gene TPM which replaces the continuous splice site usage feature in AbSplice2 with the TPM value of the gene.

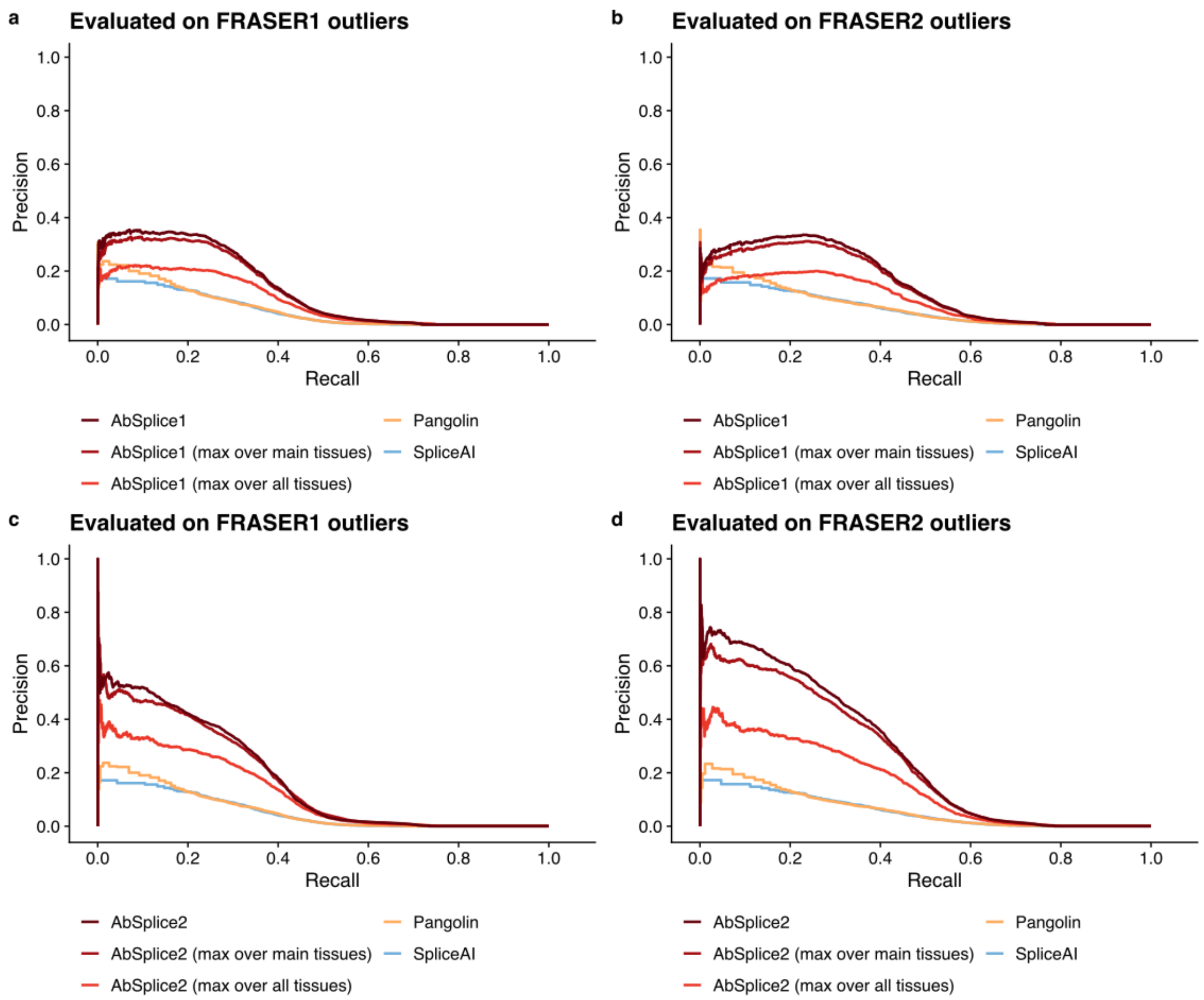


**Figure S4: Using the maximum across tissues significantly decreases performance in predicting tissue-specific aberrant splicing.** Precision–recall curves comparing the overall prediction performance on all GTEx tissues of different models evaluated on FRASER1 and FRASER2 outlier ground truth. For AbSplice models the tissue-specific scores are compared to the maximum scores across all tissues and main tissue types. **a)** SpliceAI, Pangolin and AbSplice1 models evaluated on FRASER1 ground truth. **b)** SpliceAI, Pangolin and AbSplice1 models evaluated on FRASER2 ground truth. **c)** SpliceAI, Pangolin and AbSplice2 models evaluated on FRASER1 ground truth. **d)** SpliceAI, Pangolin and AbSplice2 models evaluated on FRASER2 ground truth.

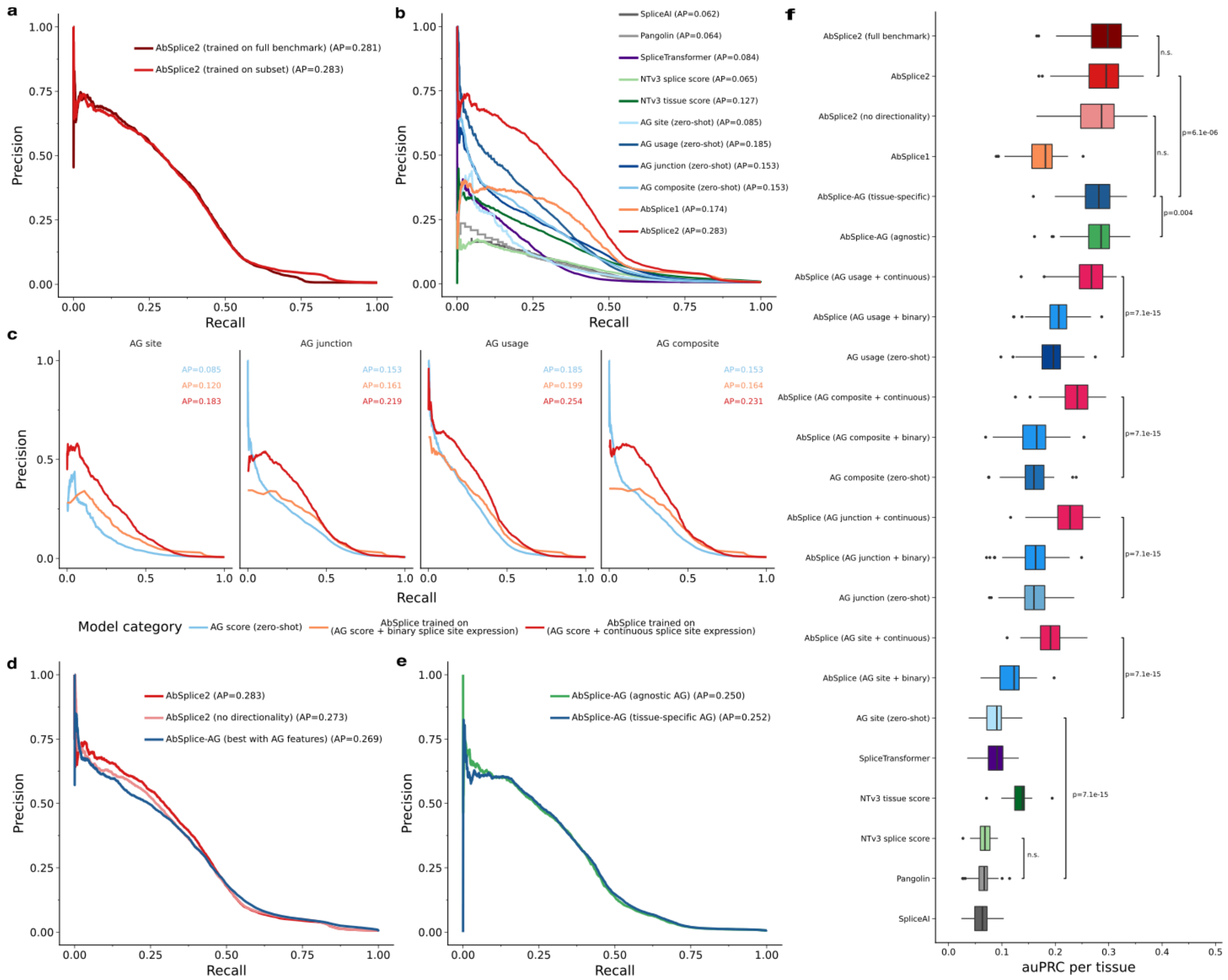


**Figure S5: Benchmarking AbSplice2 against AlphaGenome, Nucleotide Transformer v3, and SpliceTransformer. a)** auPRC of AbSplice2 trained on the full GTEx benchmark dataset vs. a model-informed subset (Methods) used for AlphaGenome (AG), Nucleotide Transformer v3 (NTv3) and SpliceTransformer variant scoring. Both AbSplice2 models achieved equivalent performance, confirming the subset is representative for training. The subset was used for all comparisons in panels **(b-f)**. **b)** auPRC of splicing predictors against GTEx splicing outliers. Sequence-based models: SpliceAI, Pangolin, AG splice scores (site, usage, junction, composite), NTv3 scores (tissue-agnostic, tissue-aware), and SpliceTransformer (tissue-specific). Trained models on the subset: AbSplice1, AbSplice2. AG splicing scores achieved the highest auPRC among standalone sequence-based models. **c)** auPRC of AG scores and AbSplice models trained with two features: one AG score type (site, usage, junction, or composite) paired with either a binary splice-site expression indicator (as in AbSplice1) or a continuous splice-site coverage feature (as in AbSplice2). **d)** auPRC comparison of the best AbSplice-AG model (Methods) against AbSplice2, as well as a version of AbSplice2 that lacks directionality information (Methods). **e)** auPRC of AbSplice models trained with tissue-agnostic AlphaGenome scores (maximum across tissues) vs. the corresponding tissue-specific AG scores, each combined with the continuous splice-site coverage feature. **f)** Per-tissue auPRC distributions (49 GTEx tissues) for all models shown in **(a-e)**. For NTv3 and SpliceTransformer predictions were only available for a subset of tissues, 34 and 33 respectively; performance was evaluated on the respective subset. Brackets show paired-two sided Wilcoxon test p-values. n.s. not significant.

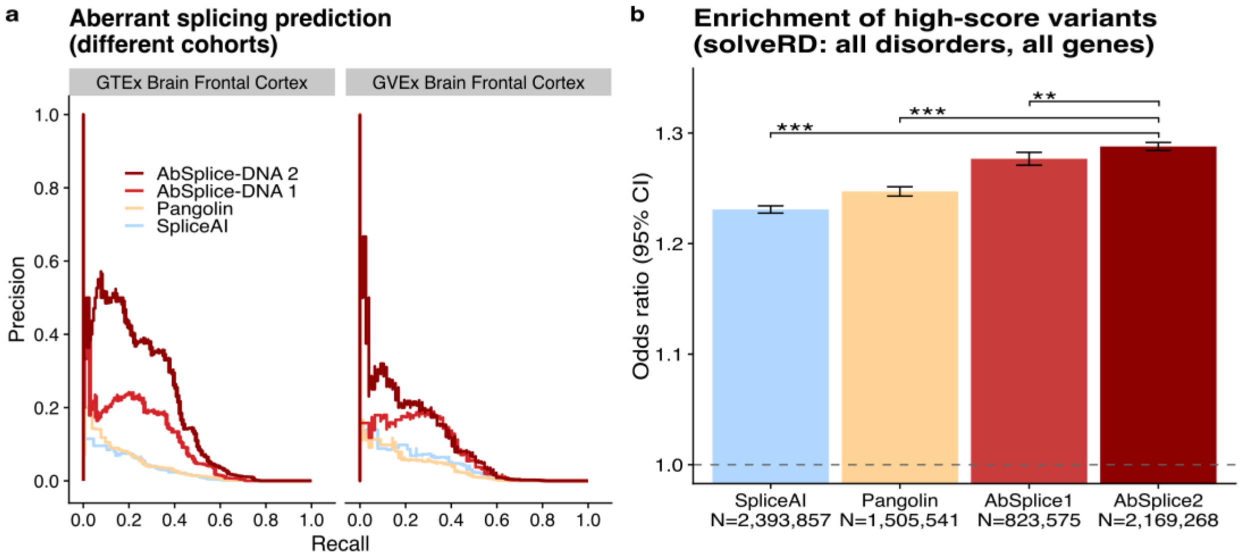


**Figure S6: Model improvements generalize to independent datasets.** Comparison of precision–recall curves for the models in (**a**) using SpliceMaps from the tissue 'Brain Frontal Cortex BA9' from the GTEx dataset to predict aberrant splicing in GTEx and the independent dataset BrainGVEx. **b)** Enrichment of high impact variants (SpliceAI: 0.8, Pangolin: 0.8, AbSplice1: 0.2, AbSplice2: 0.2) in affected individuals diagnosed with any disorder within the Solve-RD cohort. The odds ratios are computed across all genes. Error bars indicate Wald 95% confidence intervals. Asterisks mark significance levels of Wald z-tests on log odds ratios, comparing each model to the maximum AbSplice2 across tissues (*<0.05, **<0.01, ***<0.001).

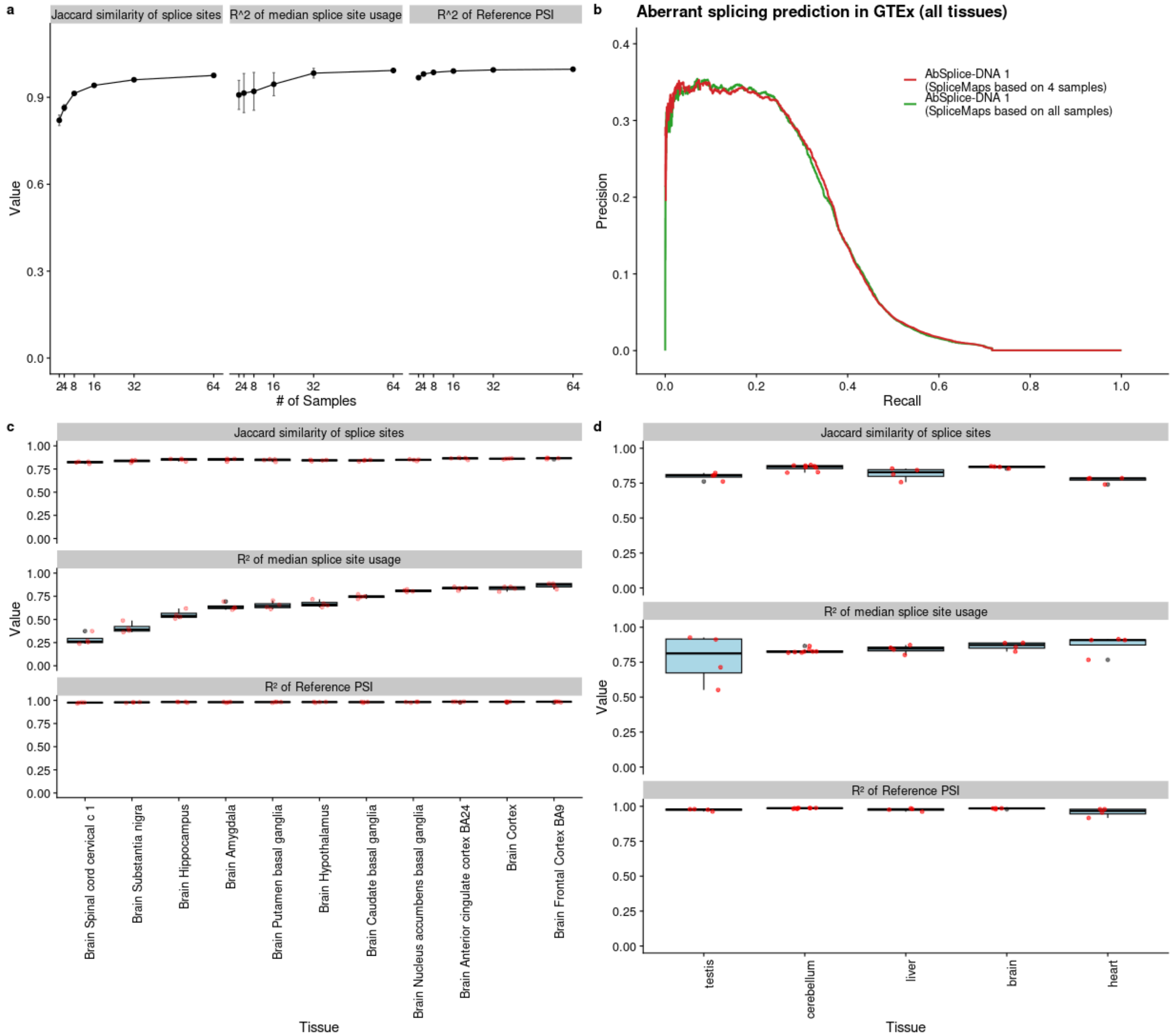


**Figure S7: Robust SpliceMaps can be constructed from even a few RNA-seq samples. a)** Comparison between SpliceMaps created from all available samples (N = 203) and different sample subsets in the GTEx tissue "Liver". Displayed metrics are the $R^2$ of $\Psi_{ref}$ (computed on shared introns in the SpliceMaps), the $R^2$ of the median splice site usage (computed on shared introns in the SpliceMaps) and the Jaccard similarity of annotated splice sites. Error bars represent one standard deviation estimated with 10 random samplings with replacement. **b)** Precision-recall curve in the GTEx benchmark dataset comparing AbSplice predictions based on SpliceMaps created from all available samples in each tissue and predictions based on SpliceMaps created from only 4 random samples in each tissue. **c)** Comparison between SpliceMaps in brain tissues from developmental stages after birth (infant, teen, adult, senior) created from RNA-seq data of Mazin et al.[25] (6<=N<=17) and GTEx SpliceMaps from all sub tissues in the brain (121<=N<=240). Displayed metrics are the $R^2$ of $\Psi_{ref}$ (computed on shared introns in the SpliceMaps), the $R^2$ of the median splice site usage (computed on shared introns in the SpliceMaps) and the Jaccard similarity of annotated splice sites. Center line, median (across developmental stages); box limits, first and third quartiles; whiskers span all data within 1.5 interquartile ranges of the lower and upper quartiles. **d)** Comparison between SpliceMaps in all available tissue types from developmental stages after birth (infant, teen, adult, senior) created from RNA-seq data of Mazin et al.[25] (3<=N<=21) and GTEx SpliceMaps from the same tissue type (121<=N<=411). For the brain and the heart in GTEx the most similar sub-tissue was chosen (brain: Brain Frontal Cortex BA9, heart: Heart Left Ventricle). Center line, median (across developmental stages); box limits, first and third quartiles; whiskers span all data within 1.5 interquartile ranges of the lower and upper quartiles.

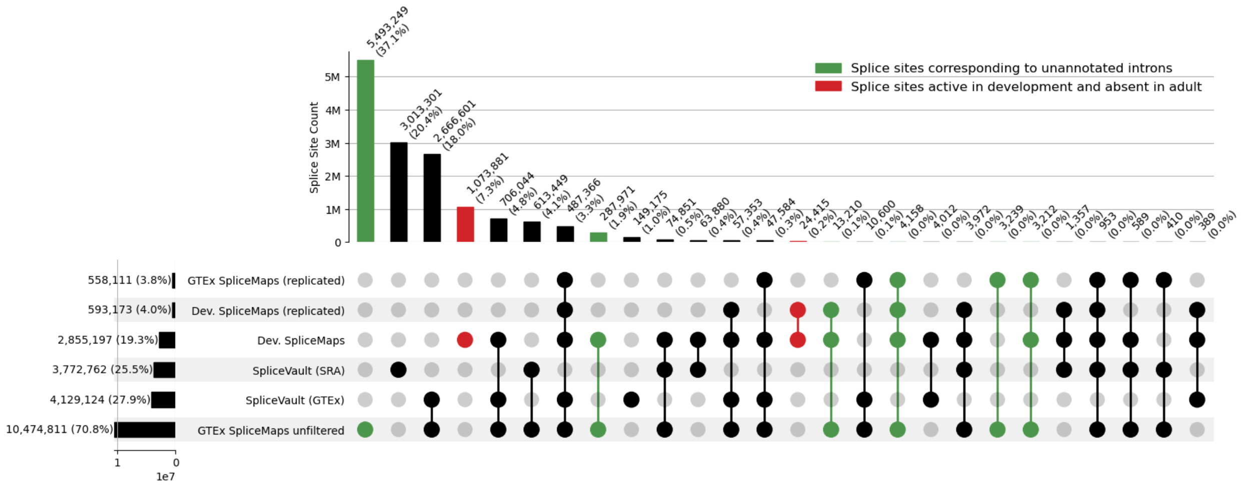


**Figure S8: Comparison of developmental SpliceMaps with population-scale splicing catalogues.** UpSet plot showing the intersection of splice sites detected in developmental SpliceMaps (both unfiltered and replicated across samples; Methods), adult GTEx SpliceMaps as used in AbSplice, adult GTEx SpliceMaps without sample reproducibility filtering ("GTEx SpliceMaps unfiltered"; Methods), and the SpliceVault 300K-RNA repository partitioned into SRA and GTEx sources. Intersection sizes are shown as counts and percentages. Green highlights: splice sites corresponding to unannotated introns (junctions where both the donor and acceptor are not present in the reference annotation). These sites are inherently absent from the SpliceVault database, which identifies mis-splicing events anchored to at least one annotated splice site. Red highlights: splice sites detected exclusively in developmental contexts. These coordinates represent developmental-specific splicing signals that are not utilized in any junctions cataloged in adult population databases, even within the high-depth SRA cohort (n=316,449 samples). The majority of detectable developmental sites (~1 million sites) represent rare, non-replicated developmental splicing events (median split-reads across individuals = 0). While SpliceMaps include these sites for maximum sensitivity, AbSplice integrates quantitative usage levels as a continuous predictive feature prioritizing replicated sites while penalizing lower-confidence signals.

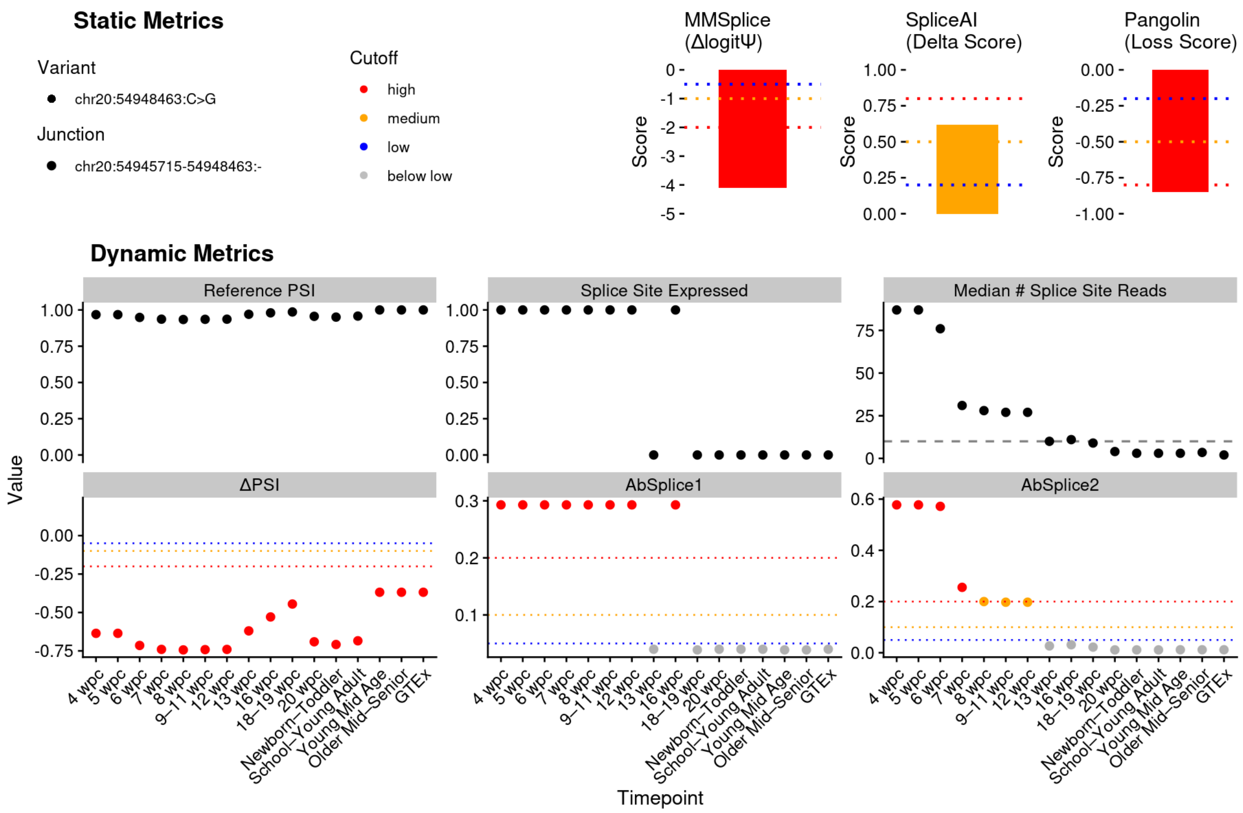


**Figure S9: AbSplice2 provides smooth predictions around splice sites with low splice site usage.** Predicted variant effects of the variant chr20:54948463:C>G (NM_198437.3:c.854+1G>C) affecting the junction chr20:54945715-54958463:- at different developmental timepoints in the brain. The deep learning models MMSplice, Pangolin and SpliceAI predict a variant effect independent of the developmental timepoint. The splice site usage of the junction changes over the course of development and is captured in the metric $\Psi_{ref}$ and the median number of split reads of the splice site annotated in the SpliceMaps of the respective timepoints. One of the input features for AbSplice1 was the binarized splice site usage with a cutoff of 10 split reads (highlighted with a dashed grey line). This binarization led to abrupt prediction shifts based on minor changes in splice site usage. AbSplice2 which made use of the non-binarized splice site usage produced much smoother prediction changes better reflecting the usage changes. Genomic coordinates are in hg19.

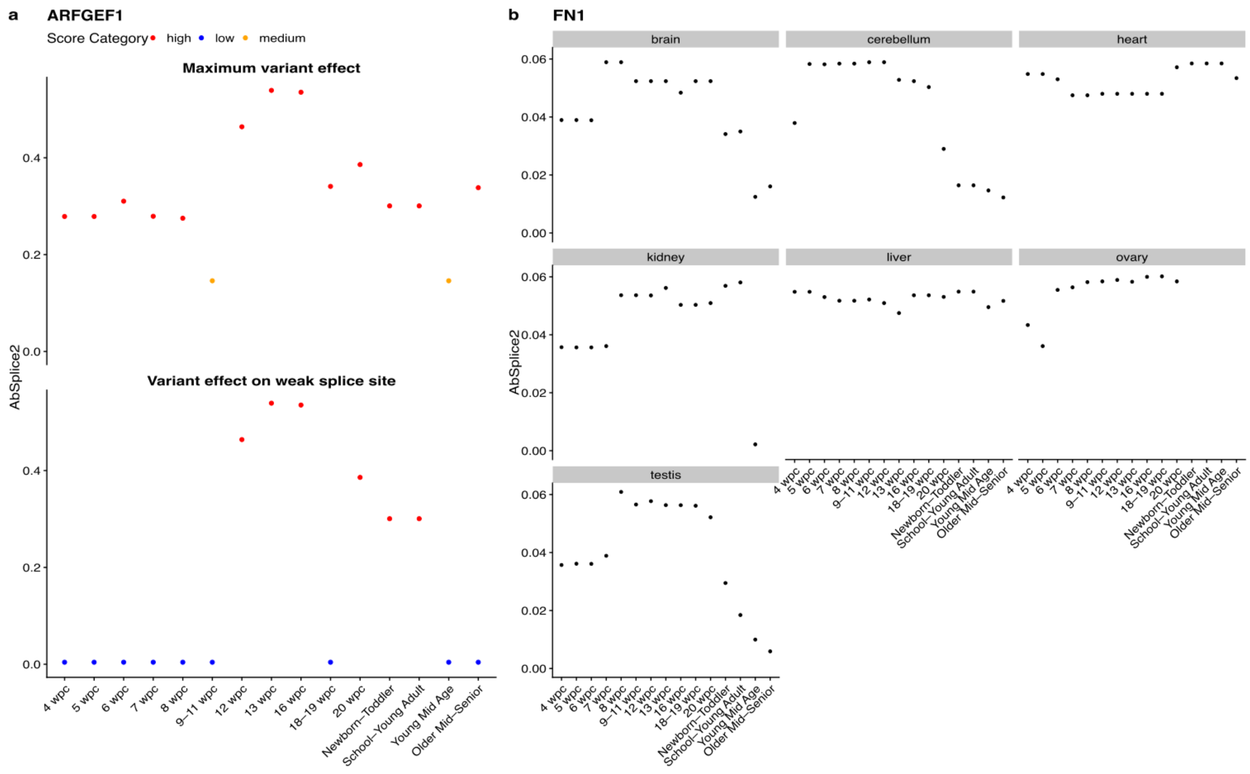


**Figure S10: Application to rare disease diagnostics. a)** The variant chr8:67203252:C>T (NM_006421.5:c.4960-1G>A) in the gene *ARFGEF1* has been identified as the causal variant in an individual recruited with intellectual disability within the NGRL. The variant activates a weak splice site within the exon causing a frameshift. Top. Maximum AbSplice2 effect predictions of the variant regarding loss of the canonical site and the gain of the weak splice site. Bottom. AbSplice2 predictions of the variant on the weak splice site. The effect is predicted to be strong in the brain between 12 and 16 weeks post-conception. **b)** The variant chr2:215372382:G>C (NM_212482.4:c.6248-7C>G) in the gene *FN1* was identified in an individual recruited to the NGRL with a diagnosis of skeletal dysplasia and developmental phenotypes. While the most relevant tissue for this phenotype was not available in our dataset, AbSplice2 predicted developmentally dynamic splicing effects across multiple tissues, with higher predicted impact during early development. This pattern supports the variant as a strong candidate, given the early clinical manifestation.

35. Aggregated Variant Calls (AggV2) - Genomics England Research Environment User

Guide. https://re-docs.genomicsengland.co.uk/aggv2/.

36. Gargano, M. A. *et al.* The Human Phenotype Ontology in 2024: phenotypes around the world. *Nucleic Acids Res.* **52**, D1333–D1346 (2024).

37. Smith, C. & Kitzman, J. O. Benchmarking splice variant prediction algorithms using massively parallel splicing assays. *Genome Biol.* **24**, 294 (2023).

38. Charles, M. *et al.* Functional impact of splicing variants in the elaboration of complex traits in cattle. *Nat. Commun.* **16**, 3893 (2025).

39. Neverov, A. & Wagner, N. AbSplice web interface. https://absplice.cmm.cit.tum.de.

45. Genomics England. The National Genomic Research Library v5.1. https://doi.org/doi:10.6084/m9.figshare.4530893/7.%25202020.

46. Martin, A. R. *et al.* PanelApp crowdsources expert knowledge to establish consensus diagnostic gene panels. *Nat. Genet.* **51**, 1560–1565 (2019).

47. Thomas, Q. *et al.* Haploinsufficiency of ARFGEF1 is associated with developmental delay, intellectual disability, and epilepsy with variable expressivity. *Genet. Med.* **23**, 1901–1911 (2021).

48. Kember, J., Gracia-Tabuenca, Z., Patel, R., Chakravarty, M. & Chai, X. J. Neurite

development varies across the hippocampus and covaries with the cellular composition of hippocampal tissue. *Commun. Biol.* **8**, 1–10 (2025).

57. Chen, K. *et al.* Self-supervised learning on millions of primary RNA sequences from 72 vertebrates improves sequence-based RNA splicing prediction. *Brief. Bioinform.* **25**, bbae163

(2024).
